# Regulatory memory and growth-coupled inheritance shape nutrient-dependent flagella number variation in *Salmonella*

**DOI:** 10.64898/2026.07.10.737737

**Authors:** Arnab Barua, Maria Giralt-Zuñiga, Marc Erhardt, Haralampos Hatzikirou

**Author notes:** Corresponding authors; e. These authors contributed equally (Arnab Barua, Maria Giralt-Zuñiga).

## Abstract

Bacteria must balance the advantage of movement against the cost of building flagella, yet how nutrient availability shapes variation in flagellar number across single cells remains unclear. Here, we combine time-resolved basal-body measurements in *Salmonella enterica* with a mechanistically constrained stochastic model of flagellar remodeling. The model separates two routes from nutrient availability to flagellar number: an RflP-dependent regulatory memory that sets a synthesis target via a latent sensing-memory variable, and a physical inheritance process in which synthesis, binomial partitioning, and division reshape the flagellar-number distribution. Coarse-graining this process yields leaky-integrator dynamics in which the mean flagellar number tracks a regulatory target, while the latent correlation follows acquisition–decay dynamics. In wild-type cells, nutrient-dependent acquisition raises the target and increases flagellar investment; in Δ*rflP* cells, loss of acquisition produces a transient overshoot that isolates the intrinsic decay (memory) timescale of the regulatory state. The fitted model predicts a held-out nutrient condition and reveals which parameter combinations are identifiable from the data. Analysis of the fitted dynamics suggests that precision is tuned primarily by sensing-dependent signal amplitude, rather than integration time, with an apparent ∼1.7-fold increase in effective wild-type noise amplitude after mean normalization, consistent with a precision cost of active regulation relative to the mutant. A model-free, information-geometric (Cramer–Rao) speed limit further shows that active remodeling approaches the statistical speed allowed by the observed distributional variability. Together, these results reveal how *Salmonella* cells couple regulatory memory with growth-dependent inheritance to record recent nutrient history in their flagellar number.

## 1 Introduction

Bacteria inhabit nearly every ecological niche on Earth, from nutrient-rich host environments to the extreme physicochemical conditions of deep-sea vents and hypersaline lakes. Their survival depends on sensing and responding to environmental fluctuations by adjusting metabolism, gene expression, and movement. Motility and chemotaxis are central to this adaptive behavior, enabling cells to explore, locate favorable habitats, and evade harmful stimuli [1, 2]. These processes integrate environmental sensing with energy expenditure, determining when it is advantageous to move versus conserve resources. Chemoreceptors detect extracellular ligands and relay information through a conserved Che phosphorylation cascade that modulates flagellar motor switching, biasing run–tumble dynamics for gradient climbing [2]. Because building and running the motility machinery is costly, cells must couple motility investment to expected payoff.

This cost–benefit trade-off is particularly consequential in facultative pathogens such as *Salmonella enterica*, where motility contributes to host colonization, tissue tropism, and immune evasion. The molecular machinery coupling nutrient state to flagellar gene expression is, as reviewed below, characterized in considerable detail. What remains missing is a quantitative, dynamical account at the level of the population: how reliably the number of flagella a cell builds encodes its nutrient environment, how the full distribution of flagellar number per cell evolves over time, and whether that distribution constitutes an information-bearing readout of regulatory state rather than a passive byproduct of an underlying switch. It is this question that we address here.

The bacterial flagellum comprises a basal body, an external hook, and a long filament that functions as a propeller [3, 4]. In *Escherichia coli* and *S. enterica*, the biogenesis of this machine requires coordinated expression of more than 60 genes in a three-layered transcriptional hierarchy [2, 5]. At the top level, the Class I *flhDC* operon encodes the master regulator FlhD_4_C_2_, which activates Class II promoters. Class II gene products include the hook–basal body, the alternative sigma factor FliA (*σ*^28^), and its anti-sigma FlgM [6]. Completion of the hook–basal body triggers FlgM secretion, releasing *σ*^28^ to drive Class III promoters encoding the filament, motor components, and chemotaxis machinery [2, 7]. This architecture enforces just-in-time assembly and couples structural progression to transcriptional control via feedback, exemplified by the FlgM–*σ*^28^ checkpoint [5, 8].

Peritrichous species typically assemble approximately 6–10 flagella per cell on average [2], yet isogenic populations are strikingly heterogeneous. Much of this heterogeneity has been described as a binary, bistable decision: transcription of *fliC* and *fliA* partitions cells into flagellar ON and OFF subpopulations [9–11], and this switch is itself nutrient-tunable at the single-cell level [12]. Such diversification may provide bet-hedging benefits during infection, as non-flagellated cells avoid flagellin-mediated immune detection while motile counterparts retain dispersal capacity, and modulation of flagellar expression during infection underscores its role as a finely tuned virulence trait [9, 10]. The number of assembled flagella per cell, however, is a distinct and graded observable: rather than a binary ON/OFF state, it reports how many hook–basal bodies a cell has committed to build, and the shape of its population distribution in isogenic *Salmonella* has been modeled explicitly [13]. It is this graded quantity, and the dynamics of its distribution, that we take as our readout.

Because producing flagella is energetically expensive, motility investment is tightly coupled to metabolic state. In enteric bacteria, a central post-transcriptional regulator of this coupling is RflP (YdiV; locus STM1344). Although related to EAL-domain phosphodiesterases, RflP lacks catalytic activity and functions as a protein–protein interaction module that modulates the activity and stability of FlhD_4_C_2_ [14, 15]. RflP binds FlhD to inhibit promoter recognition and recruits FlhD_4_C_2_ to the ClpXP protease, promoting degradation and enabling rapid shutdown of Class II transcription (Fig. 2A) [14, 16]. This dual mechanism enables a rapid and complete shutdown of the flagellar gene expression cascade. Interestingly, RflP has been shown to be involved in sensing both the metabolic state of the cell [15] and perturbations of the cell envelope [17]. In *Salmonella*, RflP is active under nutrient-poor or starvation conditions, presumably ensuring that motility is suppressed when biosynthetic resources are scarce [15]. It has been shown that the carbon storage regulator CsrA represses *rflP*, in part by binding its mRNA, linking the metabolic state of the cell to RflP expression levels [18].

Here we take a population-level view of motility investment and ask how nutrient availability shapes the number of flagella that cells build and maintain over time. Rather than tracing the full regulatory circuitry, we adopt a coarse-grained perspective suited to noisy, heterogeneous populations: if motility is costly, do bacteria adjust flagellar biogenesis in a graded, information-bearing manner as environmental conditions improve? And is that adjustment sufficiently reliable to be read out at the population level?

To address these questions, we combine single-cell measurements of flagellar basal-body number in *S. enterica* across defined nutrient regimes and time points with a dynamical model for the evolution of the flagella-number distribution. Because the regulatory task the cell faces—inferring a fluctuating nutrient state from noisy internal signals and committing biosynthetic resources accordingly—is naturally posed as decision-making under uncertainty, we adopt a Bayesian least-microenvironmental-uncertainty formalism previously developed for eukaryotic differentiation and collective cell migration [19–23]; to our knowledge, this is its first application to a prokaryotic gene-regulatory and assembly system. The model couples this RflP-dependent regulatory input to the physical dynamics of flagellar synthesis, inheritance, growth, and division, with the Bayesian construction used as a modeling premise that fixes the form of a correlation-dependent regulatory coupling. Coarse-graining gives a mean-field description in which a latent sensing-memory variable—interpretable as the accumulated RflP-dependent regulatory state of the cell —obeys acquisition–decay dynamics, while flagellar number relaxes toward the regulatory target set by that variable.

This provides a low-dimensional framework that helps us dissect regulatory function kinetically (Fig. 1). Using a Δ*rflP* mutant as a perturbation that removes the RflP-dependent acquisition branch, we separate nutrient-dependent acquisition of the regulatory signal from intrinsic decay. The acquisition-to-decay balance sets the steady-state regulatory fidelity, and in the small-correlation regime the baseline-subtracted sensing signal has a fourth-power dependence on the steady-state correlation. We quantify the precision of this encoding, estimate the effective noise cost of active regulation, and test whether the population reprograms flagella number near a model-free information-geometric speed limit [24]. Throughout, we distinguish what the data determine—kinetic rates, fidelity ratios, precision costs, and the speed bound—from what they do not, including the absolute scale of the latent correlation and the detailed parametric form of the sensing coupling. Together, these results position flagella number as a quantitative, dynamically regulated readout of nutrient-dependent regulatory state.

**Figure 1.**
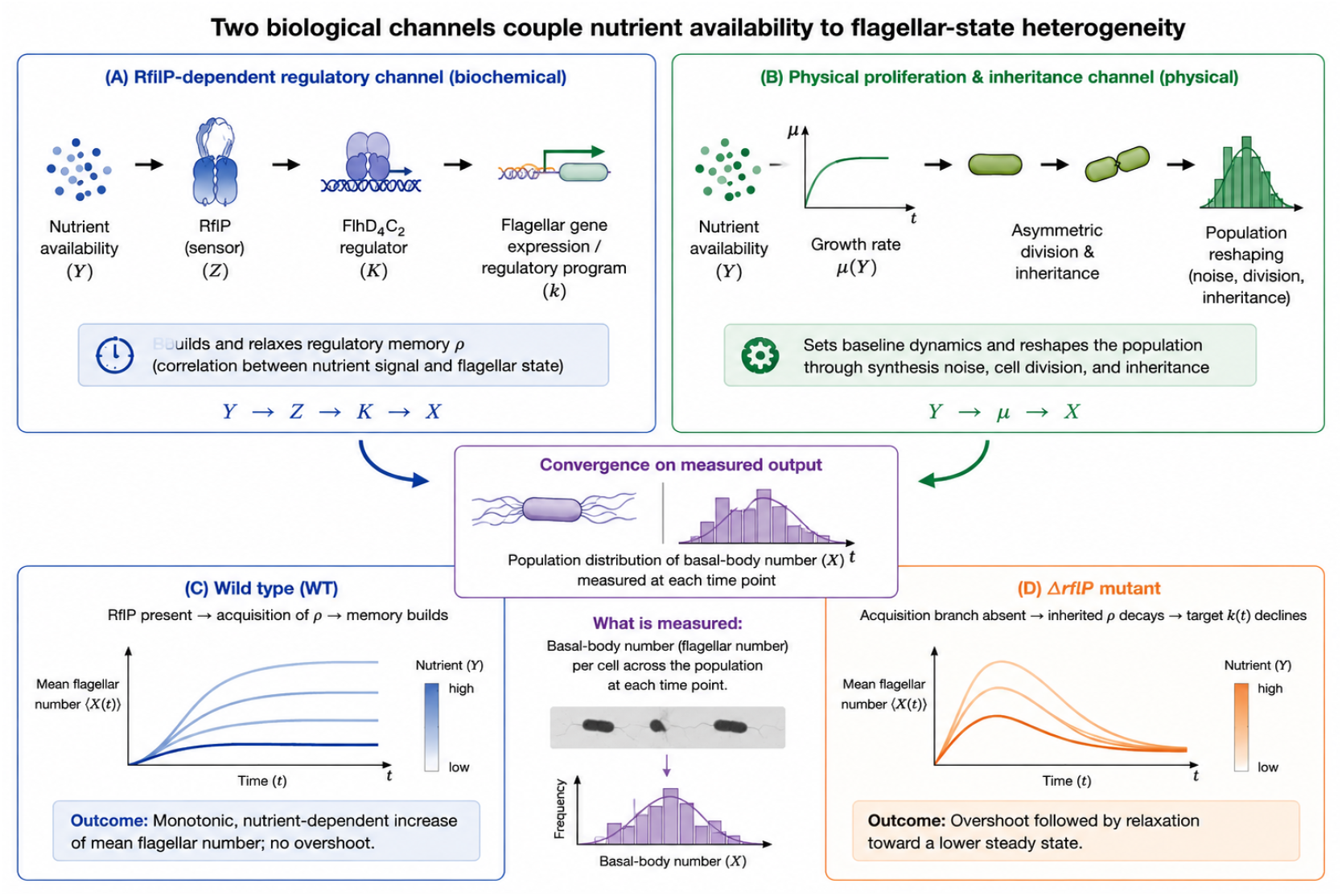
Two biological channels link nutrient availability to flagellar-state heterogeneity. Nutrients influence flagellar number through an RflP-dependent biochemical regulatory channel, summarized as *Y* → *Z K* → *X*, and through a physical proliferation and inheritance channel, summarized as *Y* → *µ* → *X*. The biochemical channel maps the nutrient-dependent regulatory state into a flagellar synthesis program, whereas the physical channel reshapes the population through synthesis noise, division, and inheritance. In the wild type both channels are active and mean flagellar number increases monotonically with nutrient; in the Δ*rflP* mutant the acquisition branch is absent, so the inherited regulatory-memory component decays and the population shows an overshoot followed by relaxation. Both channels converge on the measured population distribution of basal-body number.

## 2 Results

### 2.1 Time-resolved single-cell distributions as a population-level encoding of nutrient

To quantify the influence of nutrient availability on motility investment, we used the *S. enterica* strain EM3713, which harbors an mNeonGreen translational fusion coupled to the C-ring subunit FliG, allowing direct visualization of flagellar basal bodies (Fig. 2B). We first quantified the number of basal bodies produced at increasing nutrient concentrations by growing cells in 0.2%, 0.4%, 0.5%, and 1% yeast extract for 240 min. The distribution of basal bodies per cell increased with increasing yeast extract concentration, consistent with reduced RflP-mediated repression under high-nutrient conditions relieving the negative regulation of FlhD_4_C_2_ (Fig. 2A,C) [12, 15].

**Figure 2.**
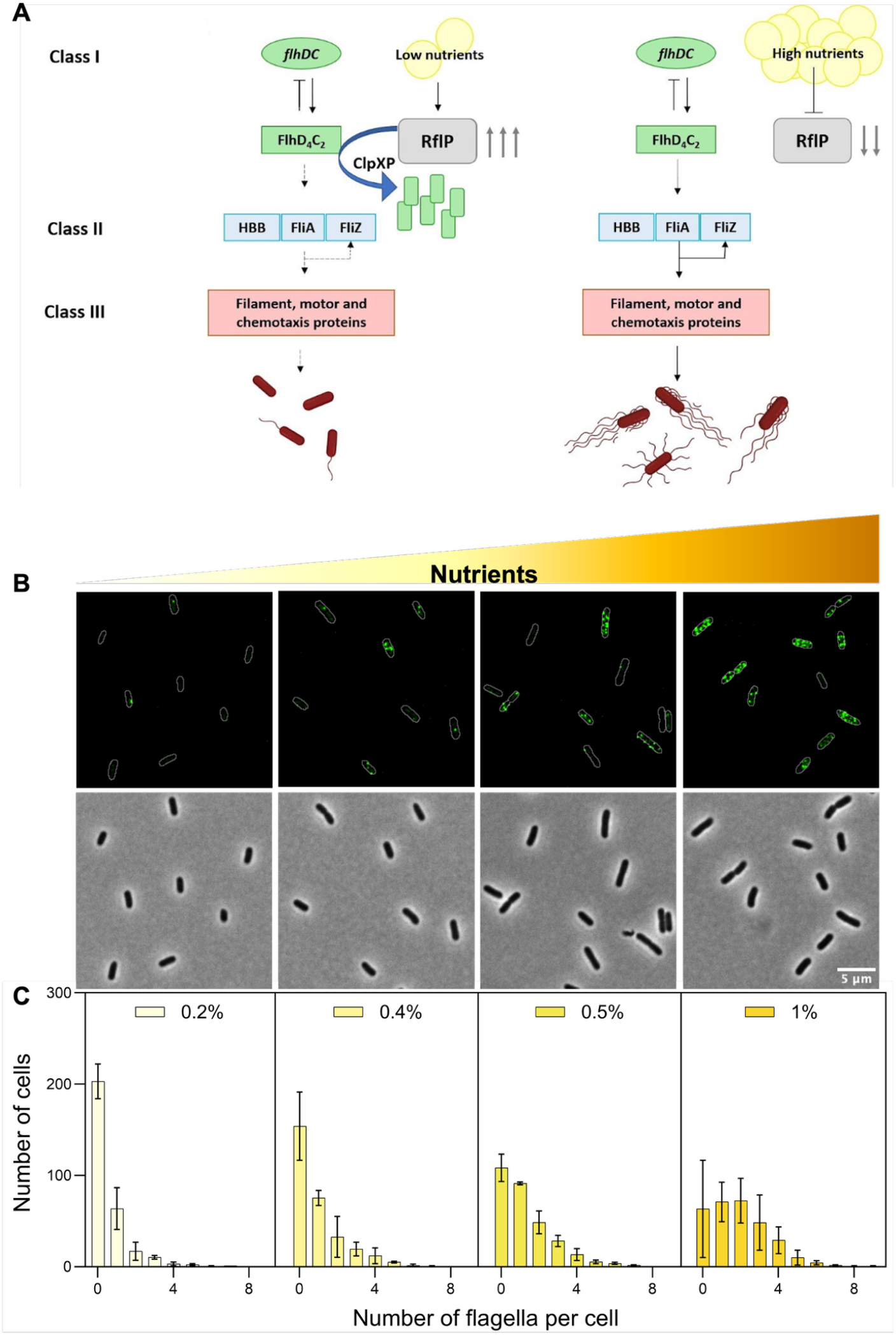
Nutrient availability tunes the fraction of flagellated and non-flagellated *Salmonella* within a population. (A) The synthesis of the flagellum involves temporal gene regulation organized into a transcriptional hierarchy of three promoter classes. Under nutrient-limiting conditions, RflP inhibits flagella synthesis by targeting FlhD_4_C_2_ for degradation by ClpXP (left panel), whereas when nutrients are abundant, RflP-mediated repression is reduced and the expression of flagellar genes is enhanced (right panel). (B) Representative microscopy images of the wild-type (WT) strain at different yeast extract concentrations, with basal bodies visualized with mNeonGreen-FliG (top panel) and phase contrast (PC) images of the bacterial cells (bottom panel). (C) Quantification of flagella (basal bodies) per cell. Measurements were performed on approx 300 cells from three independent biological replicates at 180 min.

Next, we characterized the temporal changes in flagella distribution. To ensure that cells across all nutrient conditions and time points started from a similar flagellation state, a starting culture was grown under low nutrients (0.2% yeast extract) overnight and subsequently diluted into different yeast extract concentrations. The resulting probability distributions of the flagella number are shown in Fig. 3. At low nutrient concentrations (0.2% and 0.4%), the distributions remained narrow and largely unchanged over time, indicating a modest commitment to motility. In contrast, at higher nutrient levels (0.5% and 1%), we observed a marked rightward shift in the distribution over time, reflecting a population-wide increase in flagella production. This temporal shift was especially pronounced between 0.5% and 1%, suggesting a strong sensitivity of motility investment to nutrient abundance. The data show that *S. enterica* does not express flagella at a fixed level but modulates their number over time in a nutrient-dependent manner: the distribution is largely static at low nutrient and shifts progressively rightward at high nutrient. This time-resolved, population-level response—rather than a single steady-state dose–response—is the phenomenon our model is built to explain, and it is consistent with flagellar production being a regulated, nutrient-dependent investment rather than a constitutive trait.

**Figure 3.**
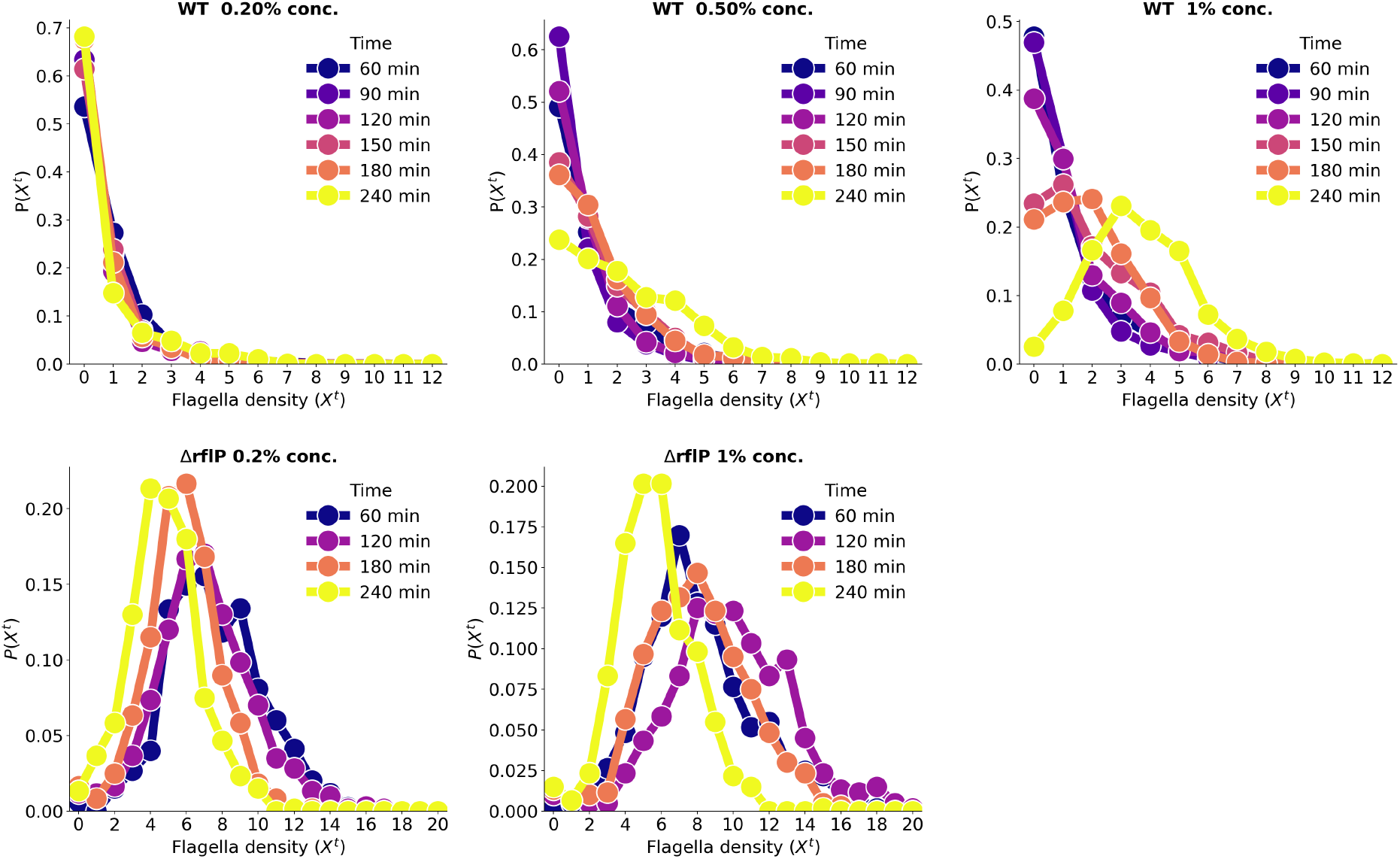
Normalized probability distribution of flagella density of WT strain and Δ*rflP* strain, averaged over 3 replicates at different time points with respect to different nutrient levels. The distributions in the top row provide the information about WT strain and the bottom row are provided for the Δ*rflP* mutant.

**Figure 4.**
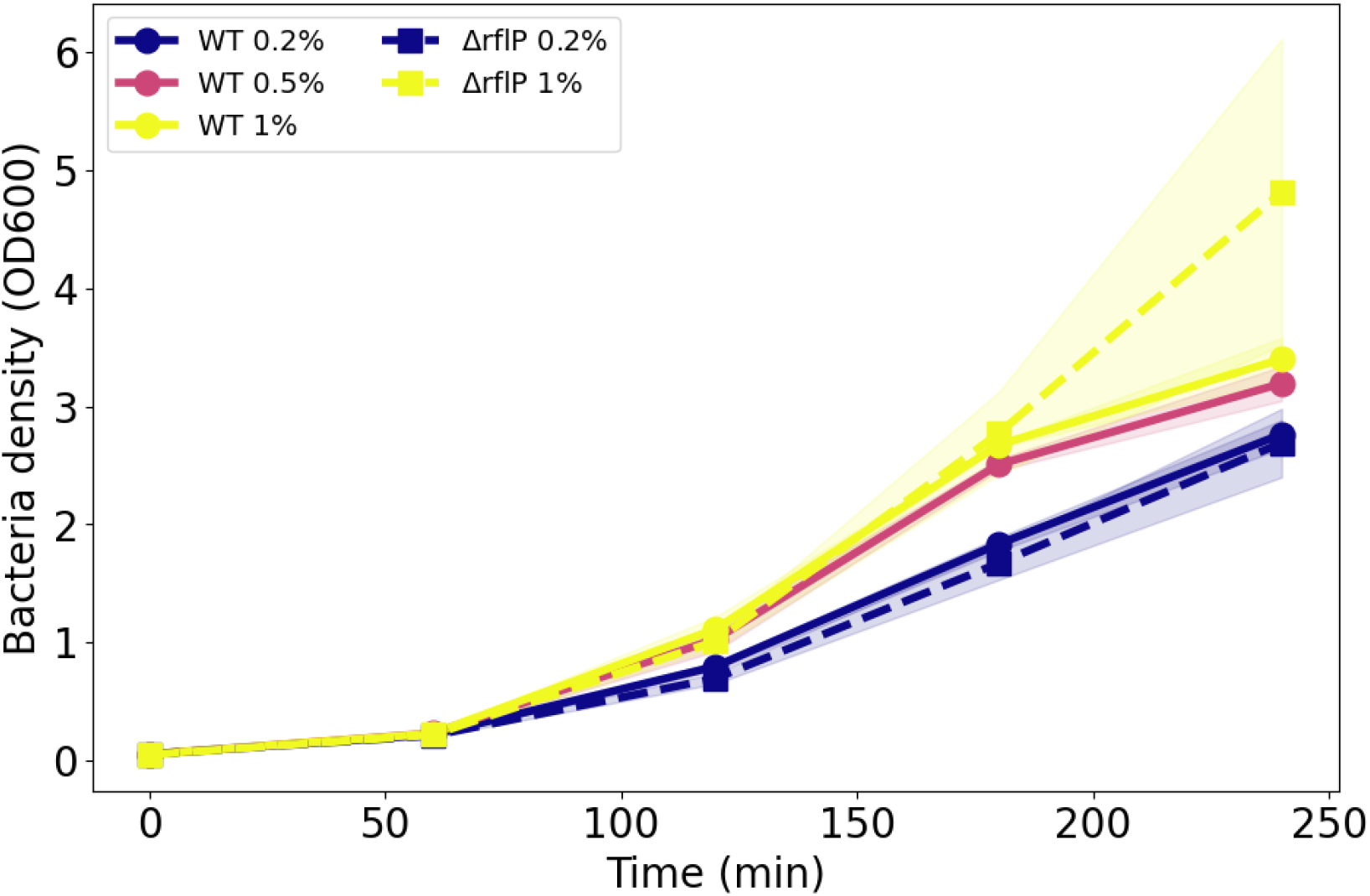
Growth rate of bacteria density for WT and Δ*rflP* strain measured at different time points at different nutrient concentration measured by OD_600_.

### 2.2 Flagellar dynamics are governed by nutrient-dependent regulation and proliferation

To capture the observed dynamics, we constructed a master equation for the probability distribution of flagellar number from two independently motivated components—an RflP-dependent regulatory input and a physical birth–partition process—and then coarse-grained it (Methods, Sections 4.5 and 4.6). The regulatory input is adapted from our earlier Bayesian sensing model [19]; here it is used as a modeling premise that supplies the correlation-dependent coupling Γ(*ρ*), rather than as a claim that these data alone prove explicit Bayesian inference by the cell. The physical component follows from Poisson synthesis of flagellar structures, binomial partitioning to daughter cells, and a load-dependent division rate. The resulting evolution equation for the distribution *P* (*x, t*) reads

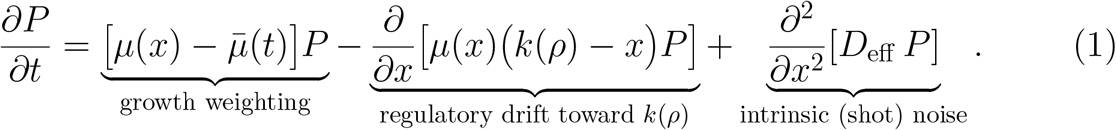

The three terms have distinct physical origins:

- **Growth weighting:** The first term reweights the distribution by relative growth. Because the division rate depends on flagellar load, *µ*(*x*) = *µ*_0_ − *ax*, subpopulations dividing faster than the population average 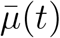 become enriched and slower ones are depleted. This is a consequence of load-dependent division and conserves total probability after normalization (Methods); it is not an externally imposed selection pressure.
- **Regulatory drift:** The second term draws the distribution toward the regulatory target *k*(*ρ*) = *k*_0_ + *c* Γ(*ρ*). Nutrient dependence enters through the latent sensing-memory variable *ρ* and the coupling *c*. Larger *ρ* raises the target, and the physical flagellar count relaxes toward it.
- **Intrinsic noise:** The final term is diffusion in flagellar-number space arising from Poisson–binomial stochasticity of synthesis and partition, with effective coefficient *D*_eff_.

Equation (1) is therefore a coarse-grained representation of a specified birth– partition process driven by an RflP-dependent regulatory input.

### 2.3 Flagellar encoding as a leaky integrator

Coarse-graining the master equation (1) yields a closed description in which the mean flagellar number relaxes toward the regulatory target *k*(*ρ*) = *k*_0_ + *c* Γ(*ρ*), while the latent sensing-memory variable *ρ*(*t*) obeys linear acquisition–decay dynamics,

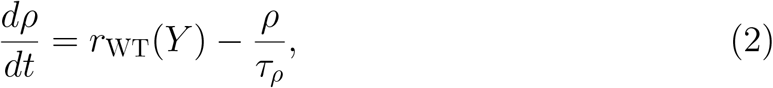

with acquisition rate *r*_WT_(*Y* ) and intrinsic decay time *τ*_*ρ*_. The solutions are

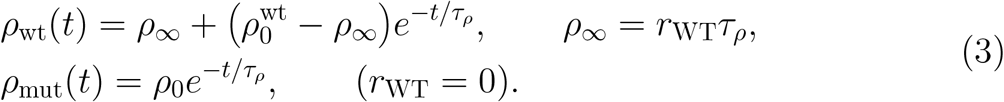

Thus the *sensing-memory variable* is the leaky integrator: nutrient-dependent acquisition builds *ρ*, whereas intrinsic decay erodes it. Flagellar number is the slower physical readout that tracks the moving target *k*(*ρ*(*t*)). In the wild type, *ρ* relaxes toward *ρ*_∞_ = *r*_WT_*τ*_*ρ*_; in the Δ*rflP* mutant the acquisition source is absent, so the inherited correlation decays toward zero (Fig. 6). This distinction explains the qualitative behavior in Fig. 5: wild-type cells build a rising target, whereas mutant cells relax back toward baseline after an initial overshoot. Here *µ*_0_ is the baseline division/growth parameter, *c* is the sensing coupling, and 1*/τ*_*ρ*_ is the intrinsic decay rate.

**Figure 5.**
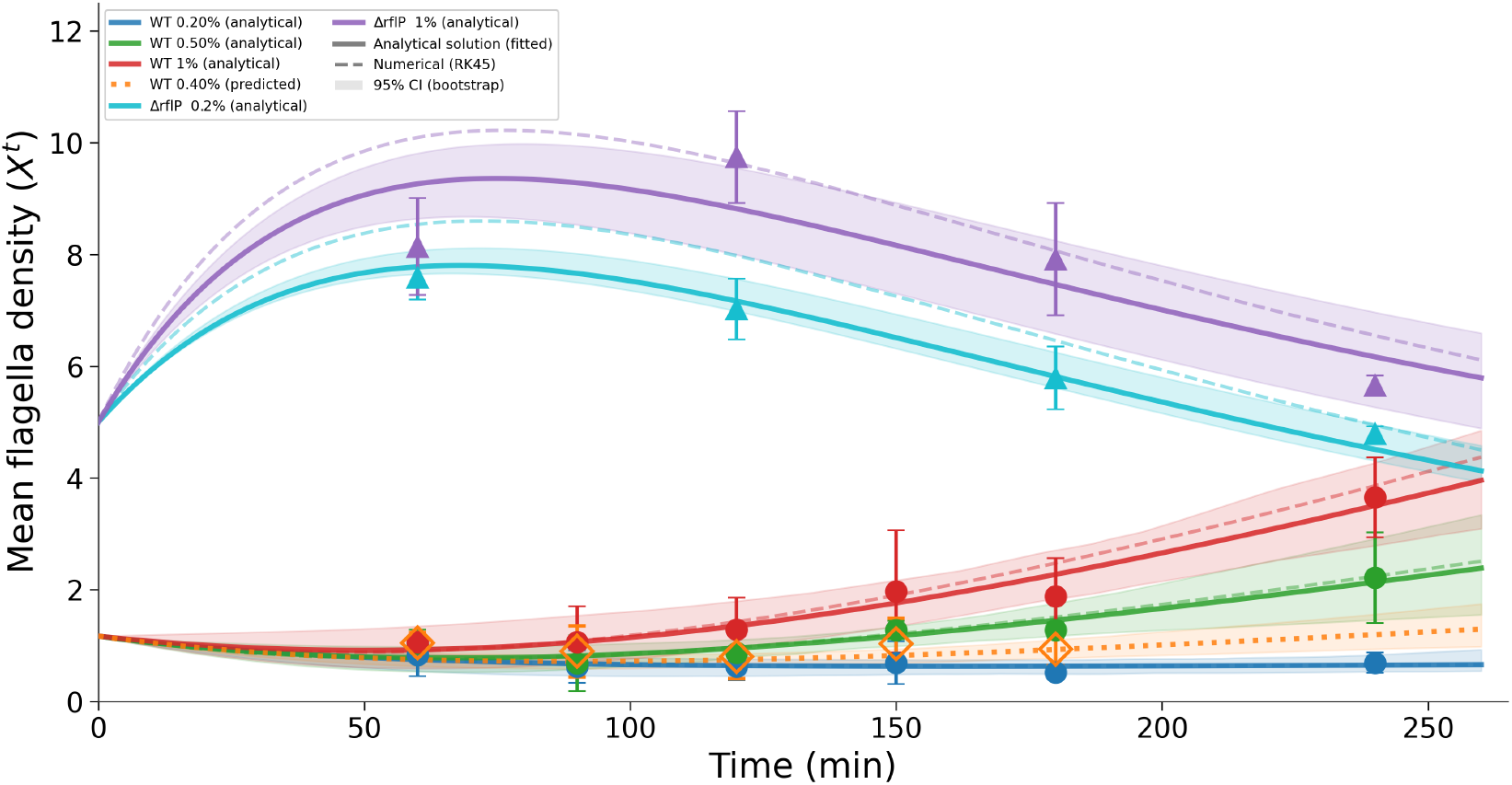
Mean-field fits to the flagellar-number trajectories. Points show experimental mean basal-body counts *X*^*t*^ for the wild type (0.2%, 0.5%, 1%; circles, rising monotonically with nutrient) and the Δ*rflP* mutant (0.2%, 1%; triangles, overshooting and relaxing). Solid lines show the fitted closed-form (small-correlation) mean-field solutions; the dotted line shows the held-out WT 0.4% prediction (open diamonds: 0.4% data, not used in fitting). Numerical integration of the full mean-field model (RK45) is overlaid as a consistency check, and shaded bands are 95% confidence intervals.

**Figure 6.**
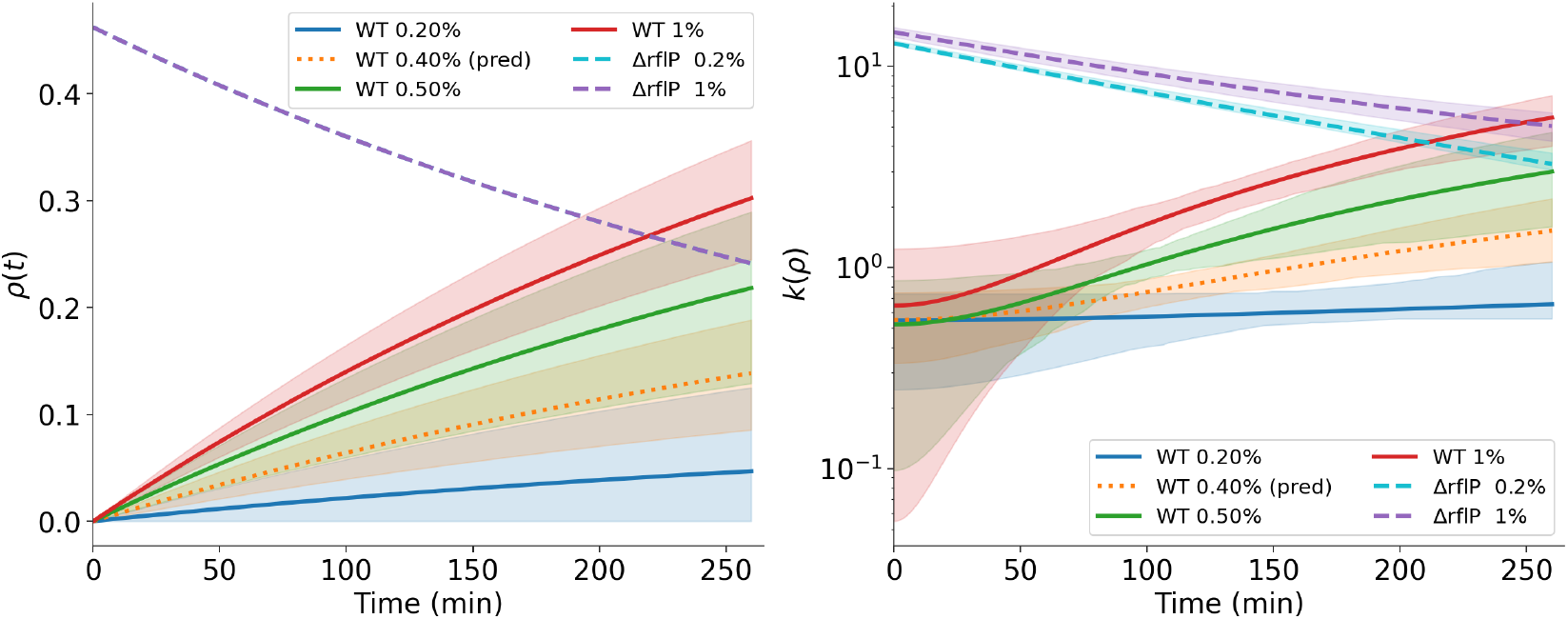
Latent sensing correlation *ρ*(*t*) across nutrient conditions and time (left), increasing in the wild type and decaying in the Δ*rflP* mutant, with the corresponding regulatory target *k*(*ρ*) = *k*_0_ + *c* Γ(*ρ*) (right, log scale).

#### 2.3.1 The Δ*rflP* mutant overshoot isolates the intrinsic decay rate

Because the mutant lacks the acquisition source (*r*_WT_(*Y* ) = 0), its dynamics depend on the intrinsic decay time *τ*_*ρ*_. The mutant therefore removes the RflP-dependent acquisition term, allowing the decay of the inherited effective regulatory state to be constrained, as shown in Fig. 6. A joint fit of both mutant nutrient conditions reveals a common, nutrient-independent decay: the mean flagellar number rises to a transient maximum and then relaxes, the signature of a correlation that starts above its (zero) steady state and decays. From this global fit we extract a characteristic memory lifetime around 400 min, corresponding to the decay timescale *τ*_*ρ*_; the precise value depends on the gauge convention for the latent correlation and is reported as a range (Methods, Section 4.10). The overshoot peak occurs at a nearly constant time of ≈ 70 min (for Δ*rflP* 0.2%) and ≈ 78 min (for Δ*rflP* 1%), a gauge-invariant feature indicating that the transient memory dynamics are set by the inherited initial correlation and the intrinsic decay, not by the nutrient level. The fitted correlation functions decay smoothly after the peak, consistent with the interpretation that, without an active acquisition source, correlations cannot be maintained and flagellar production returns toward its baseline.

#### 2.3.2 The acquisition rate increases with nutrient

The acquisition rate *r*_WT_(*Y* ) measures how quickly nutrient sensing builds the correlation that drives flagellar production. Fitting the coupled mean-field model across nutrient conditions, we find that the baseline division/growth parameter *µ*_0_ varies little across wild-type conditions, while *r*_WT_(*Y* ) increases steadily with nutrient concentration (Fig. 7B,C). The steady-state correlation *ρ*_∞_ = *r*_WT_*τ*_*ρ*_ therefore also rises with nutrient, and does so smoothly rather than in a switch-like manner—evidence that *S. enterica* tunes flagellar investment in a graded fashion across the nutrient range, rather than through an all-or-none ON/OFF response. The wild type acquires correlation over time while the mutant loses it; the absolute initial correlation *ρ*_0_ is not determined by the data and our conclusions do not depend on the specific methods (Methods, Section 4.10). The fitted *r*_WT_(*Y* ) and the resulting *ρ*_∞_ both increase with nutrient but sub-saturate at high nutrient. By contrast, the Δ*rflP* mutant lacks this nutrient-responsive acquisition (*r*_WT_ = 0), so its correlation cannot build and instead decays at the intrinsic timescale. The comparison makes explicit that nutrient sensing controls the acquisition of correlation, while the mutant overshoot exposes the underlying decay.

**Figure 7.**
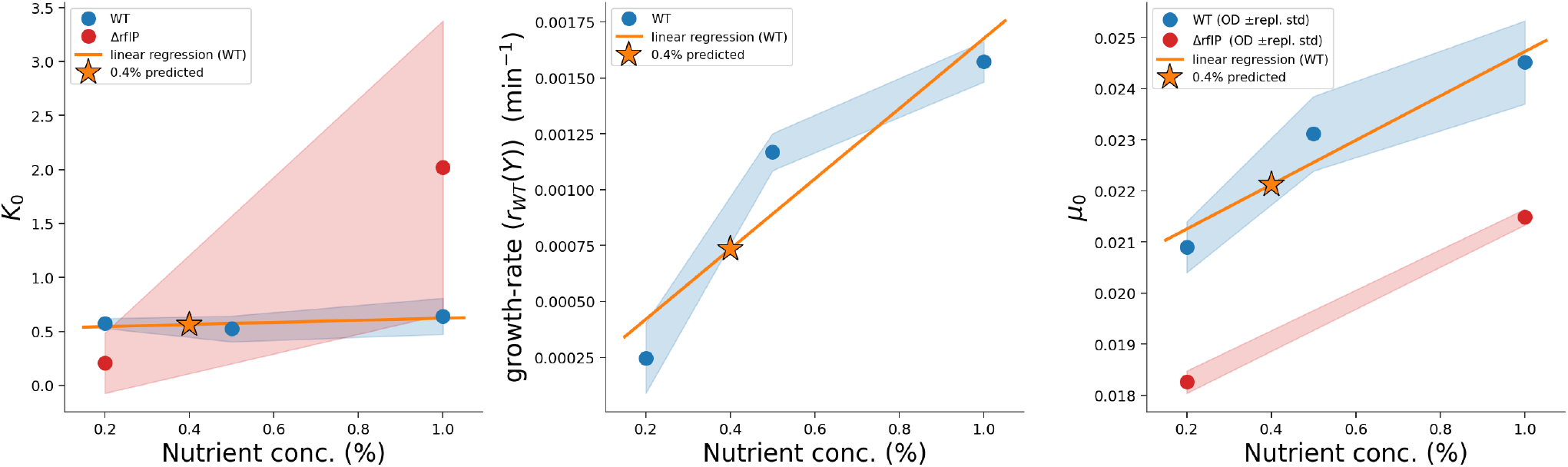
Fitted parameters versus nutrient concentration: (left) baseline target *k*_0_, approximately flat for the wild type and rising for the mutant; (middle) acquisition rate *r*_WT_ = *ρ*_∞_ */τ*_*ρ*_, increasing and sub-saturating with nutrient; (right) growth rate *µ*_0_, varying over different nutrient concentration. Orange stars/diamonds mark the held-out 0.4% prediction, which falls on the fitted trends. Shaded bands are 95% bootstrap confidence intervals.

We fix the biosynthetic cost at *a* = 2.65 × 10^−4^ and the sensing coupling at *c* = 100 (only the gauge-invariant product *c* Γ(*ρ*) is identifiable; Methods, Section 4.10). The initial correlation *ρ*_0_ is a weakly constrained (sloppy) parameter that the data do not determine (Section 4.10). Our conclusions depend only on the gauge-invariant kinetic quantities and not on its value. The fitted mean trajectories, the latent correlation dynamics, the held-out 0.4% prediction, and the nutrient dependence of the parameters are shown together in Fig. 7; the goodness-of-fit metrics are summarized in Tables 2 and 3.

**Table 1.** Globally shared parameters for the hybrid model. The normalised objective value is 0.044204.

| Parameter | Description | Value | Units |
| --- | --- | --- | --- |
| $\rho_{0,WT}$ | WT initial latent correlation | 0.000100 | — |
| $\rho_{0,mut}$ | Mutant initial latent regulatory-memory coordinate | 0.462059 | — |
| $\tau_\rho$ | Latent regulatory-memory decay time | 400.000 | min |
| $X_{0,WT}$ | WT initial condition | 1.16899 | — |
| $X_{0,mut}$ | Mutant initial condition | 5.00000 | — |

**Table 2.**
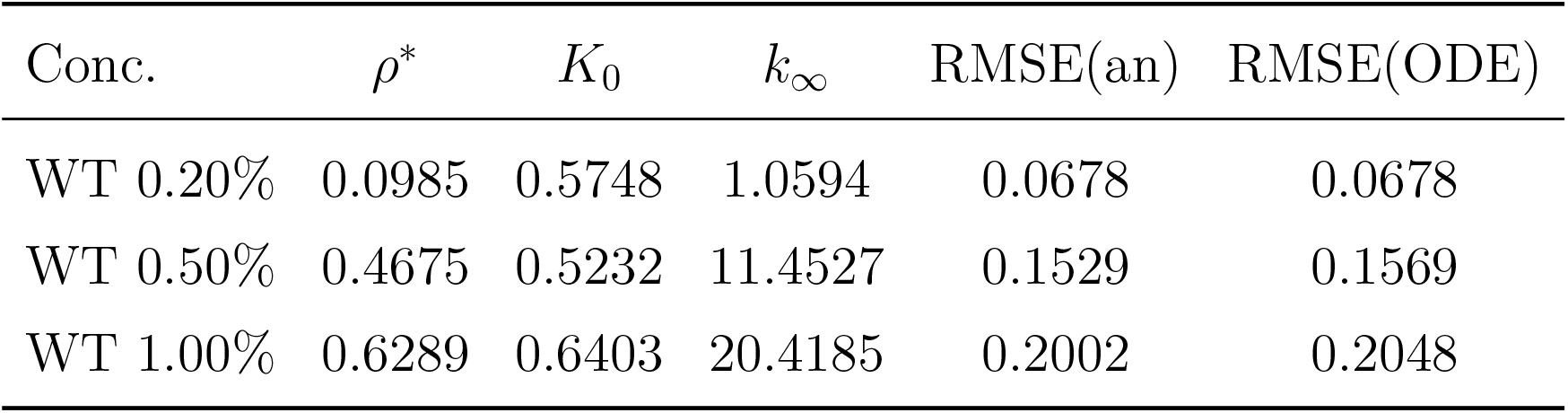
Wild-type hybrid model fit results across nutrient conditions. RMSE(an) is computed from the closed-form small-correlation analytical solution; RMSE(ODE) from numerical integration of the mean-field equations.

**Table 3.**
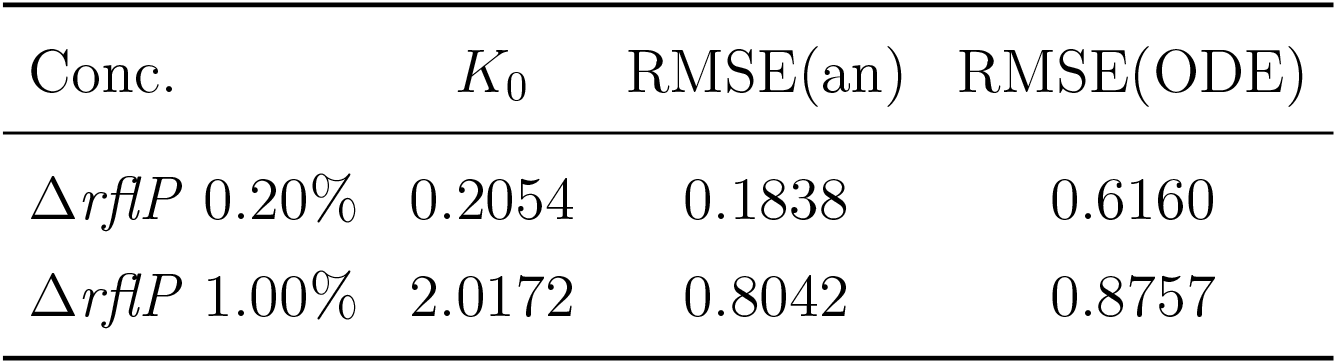
Δ*rflP* mutant hybrid model fit results across nutrient conditions. RMSE(an) is computed from the closed-form small-correlation analytical solution; RMSE(ODE) from numerical integration of the mean-field equations.

**Table 4.**
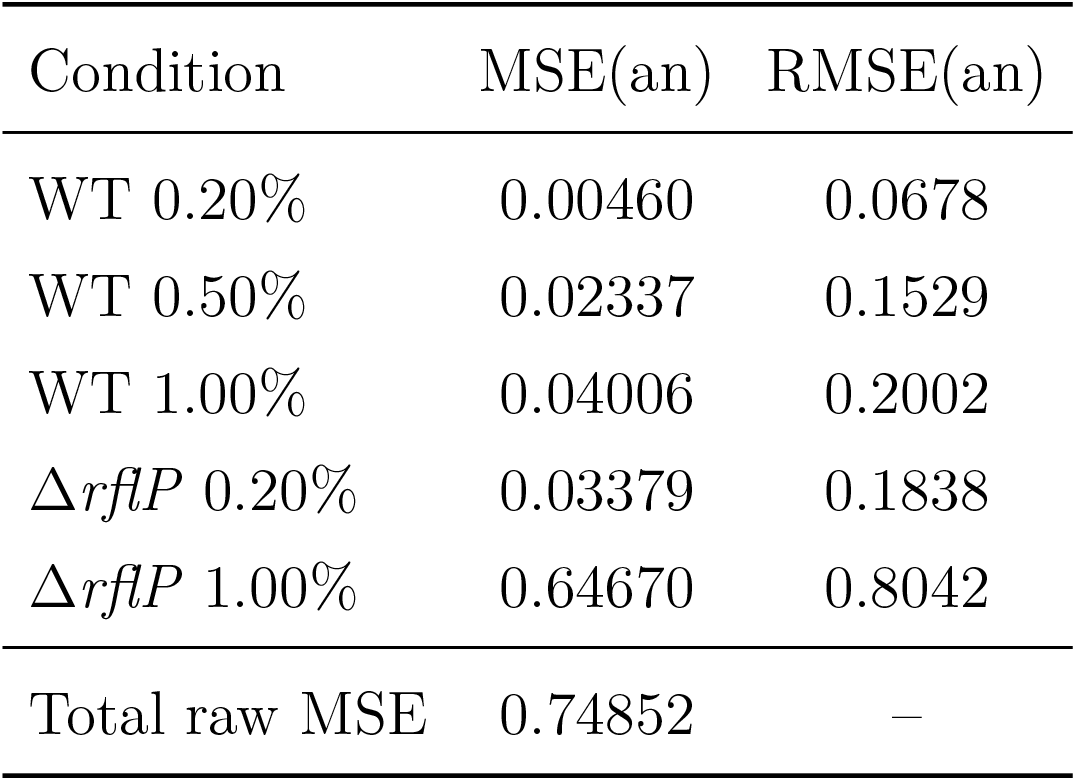
Per-condition analytical mean-squared error and root-mean-squared error for the hybrid model.

**Table 5.**
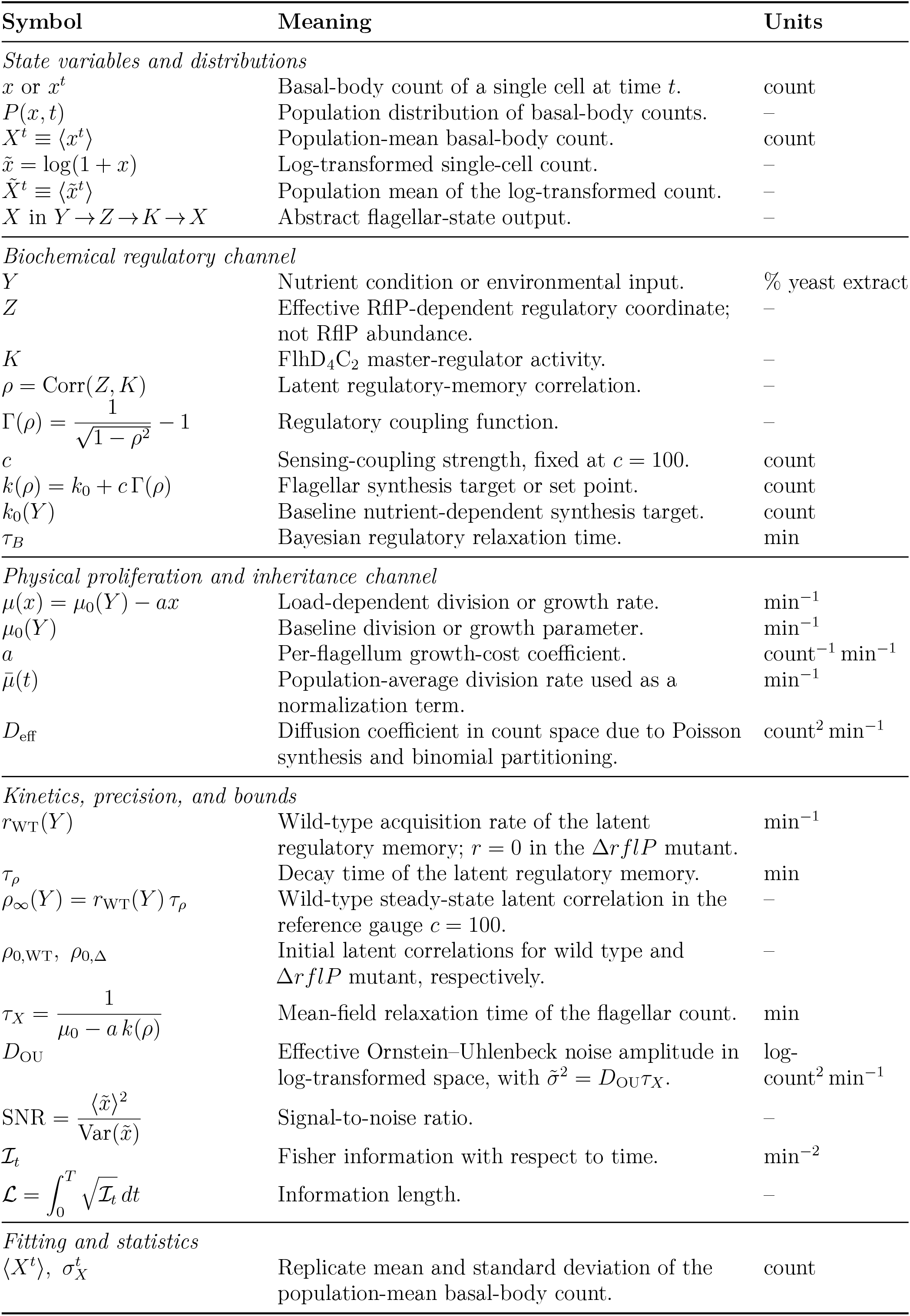
Mathematical notation summary.

Across wild-type conditions the residuals are comparable to the replicate variability. With only three replicates in hand they more likely reflect conservative uncertainty estimates or correlations among time points. The mutant fits, summarized by MSE and RMSE, capture the overshoot and relaxation. Together, the wild-type and mutant fits support a graded, homeostatic regulation of flagella number: low nutrient yields a low steady-state number (an energy-saving regime) and high nutrient a higher steady-state number, with a persistent baseline maintained throughout. We quantify the precision of this tuning below.

### 2.4 Encoding precision is tuned through signal amplitude, not

Producing and operating flagella is metabolically costly, so the precision with which a population aligns flagellar number to nutrient is a useful measure of regulatory performance. We quantify precision using the signal-to-noise ratio of the log-transformed flagellar distribution, 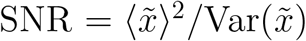 (Eq. 39). When the dynamics are linearized near an operating point, the fluctuations obey an Ornstein–Uhlenbeck approximation with stationary variance

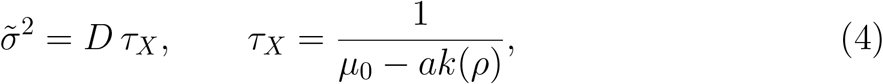

where *τ*_*X*_ is the mean-field relaxation time of the flagellar count and *D* is an effective strain- and condition-dependent noise amplitude in log-transformed space. This is a nonequilibrium variance–relaxation relation for the fitted stochastic dynamics, not an equilibrium fluctuation–dissipation theorem. The *D* should therefore not be read as a thermodynamic temperature or dissipation rate.

The SNR increases with nutrient concentration and rises over time in the wild type, whereas in the Δ*rflP* mutant it decreases over time at both nutrient levels (Fig. 8b), consistent with acquisition of the regulatory correlation in the wild type and loss of that correlation in the mutant. Two features are notable.

**Figure 8.**
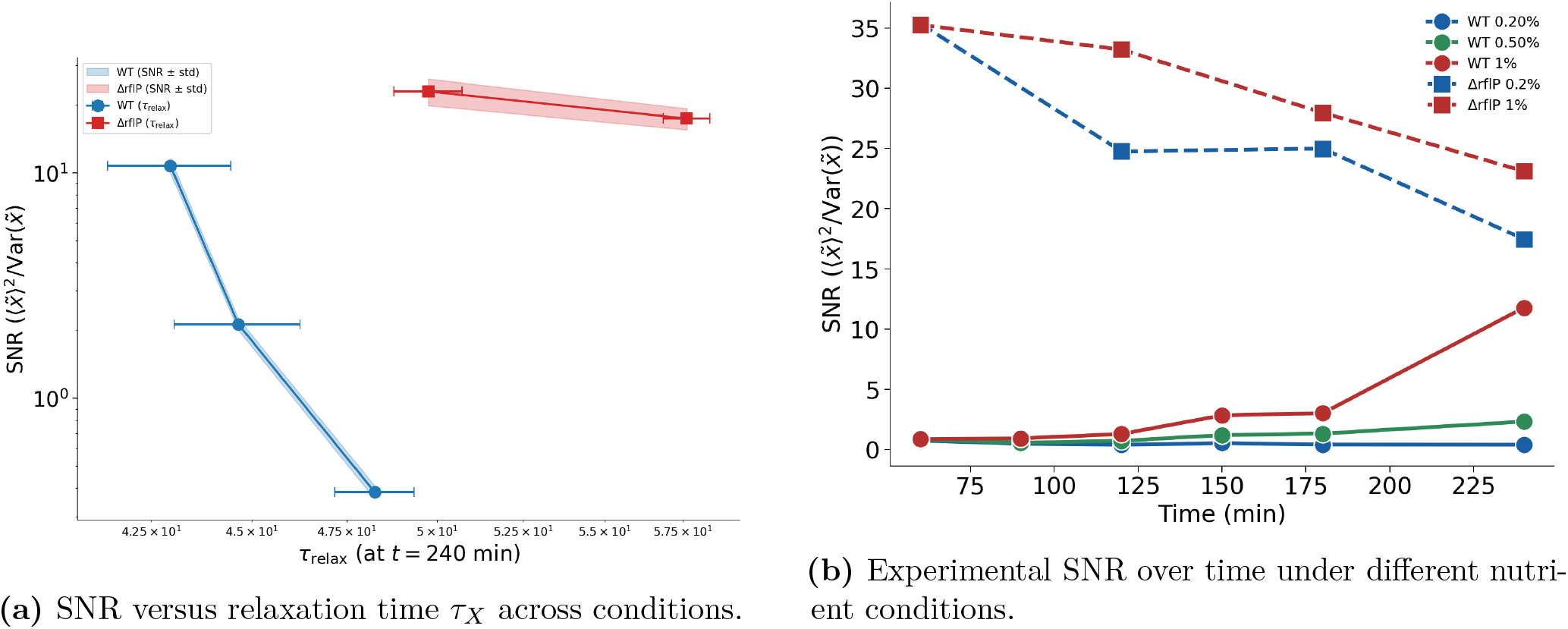
(a) Signal-to-noise ratio (SNR) versus the relaxation time *τ*_*X*_ across conditions: SNR varies by nearly two orders of magnitude while *τ*_*X*_ varies by less than a factor of two, so precision is tuned through signal amplitude rather than bandwidth. (b) Experimental SNR over time for different nutrient conditions, increasing in the wild type and decreasing in the mutant.

First, the observed precision changes mainly through signal amplitude rather than through bandwidth. Across conditions, *τ*_*X*_ varies much less than the SNR (Fig. 8a), so a simple bandwidth-limited prediction SNR ∝ 1*/τ*_*X*_ does not explain the data. In the small-correlation regime, the sensing-dependent excess target is

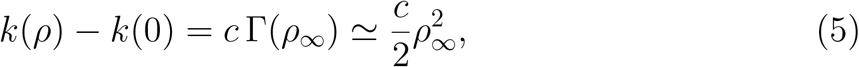

and the corresponding *sensing-dependent* signal power scales as 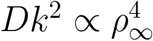 . This fourth-power statement applies to the baseline-subtracted regulatory si∞gnal; it should not be conflated with the raw 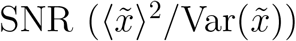 when the baseline *k*_0_ is appreciable. This is the opposite of the canonical Berg–Purcell [25] route to sensing precision, in which precision improves with the available integration time; here the relaxation time is nearly fixed across conditions and precision is instead set by the amplitude of the sensing-dependent signal.

Second, active regulation appears to carry an effective precision cost. We estimate the effective noise amplitude *D* from the variance–relaxation relation 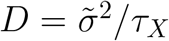, using the variance 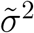 of the log-transformed flagellar count 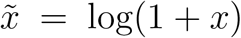. Working in log space makes the comparison mean-normalized, so it reflects relative cell-to-cell noise rather than absolute spread. On this basis the wild type carries the larger effective noise amplitude, *D*_*W T*_ */D*_Δ*rflP*_ ≈ 1.7 (range ∼ 1.5–2 across conditions and the two mutant replicates). We interpret this as an effective noise cost of maintaining a driven, nutrient-responsive regulatory signal, not as a calibrated thermodynamic dissipation. Because the inferred ratio is sensitive to the mutant growth parameter and to replicate uncertainty, we report it as an effective strain-dependent noise scale rather than a precisely calibrated cost. We emphasize that the direction depends on this normalization: in *absolute* (linear-space) terms the mutant has the higher variance, simply by virtue of its larger mean flagellar number, and the regulation-attributable excess noise in the wild type emerges only once the mean is normalized out.

### 2.5 Functional combinations are identifiable, while the latent sensing scale is not

The full model has 16 parameters across the two strains, but not all combinations are equally constrained by the data. To establish that the biological conclusions rest on well-determined quantities, we analyzed the curvature of the fit objective at the best-fit point through the Fisher-information matrix (FIM), evaluated in log-parameter space (Methods, Section 4.10). Uncertainty bands and parameter intervals are generated as described in Methods, whereas the identifiability structure is characterized through the Fisher information rather than by resampling. The FIM eigenvalues measure how stiffly the fit objective rises along the corresponding eigenvector directions in parameter space.

The eigenvalue spectrum spans roughly 10.6 orders of magnitude (Fig. 9, left), the hallmark of a “sloppy” model in which a few stiff parameter combinations dominate the fit while the remainder are weakly constrained. Please note that, for the FIM analysis we fitted separately the 0.4% nutrient concentration to see the whole picture. A shoulder near the eleventh eigenvalue separates a stiff band of eleven well-constrained directions from a steeply collapsing sloppy tail. Crucially, the composition of these directions is informative rather than arbitrary (Fig. 9, right). The stiffest eigenvectors load on the mutant initial correlation and the intrinsic decay rate 1*/τ*_*ρ*_ (together fixing the overshoot), and on the wild-type acquisition rates *r*_WT_(*Y* )—equivalently on the steady-state correlation *ρ*_∞_ = *r*_WT_*τ*_*ρ*_ that sets the encoding fidelity. Because the sensing coupling is fixed at the reference value *c* = 100 (only the product *c* Γ(*ρ*) is identifiable), *c* is not itself a coordinate of this FIM; the sloppiest eigenvectors instead load on the latent-correlation amplitudes—the wild-type initial correlation 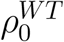 and the per-condition steady-state correlations—and on the unobserved initial conditions, which are the practical image of the structural *c*–*ρ* gauge.

**Figure 9.**
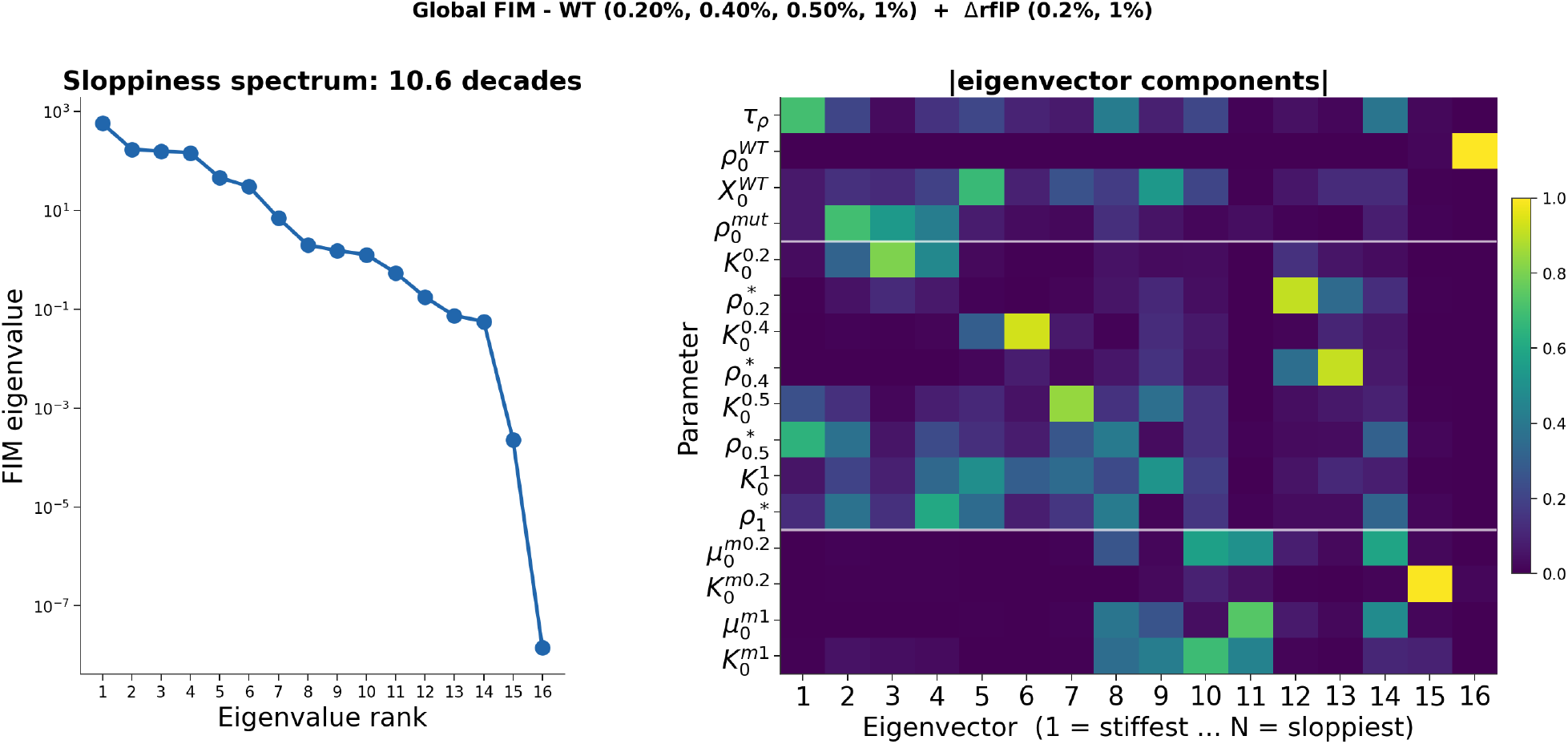
Identifiability and sloppiness analysis. (Left) Eigenvalue spectrum of the Fisher information matrix at the best fit, in log-parameter space, spanning ∼ 10.6 decades; a shoulder near the eleventh eigenvalue separates stiff (well-constrained) from sloppy (weakly constrained) directions. (Right) Absolute components of the corresponding eigenvectors, with the 16 model parameters (rows) ordered against eigenvectors from stiffest (left) to sloppiest (right). Stiff directions load on the decay rate, the mutant initial correlation, and the acquisition rates *r*_WT_(*Y* ); sloppy directions load on the latent-correlation amplitudes and unobserved initial conditions (the practical image of the *c*–*ρ* gauge; the sensing coupling *c* is fixed at the reference value and is not a FIM coordinate), confirming that the data constrain the functional kinetic quantities while leaving the gauge freedoms and unobserved initial conditions undetermined.

The data therefore pin down precisely the combinations the theory identifies as functional—the kinetic rates and the fidelity ratio—and leave unconstrained the latent-correlation amplitudes and the unobserved initial states. The absolute scale of *ρ* and the acquisition rates are gauge-dependent, since only the product *c* Γ(*ρ*) enters; the numerical estimate *τ*_*ρ*_ ≈ 400 min (reference parameterization) is not itself a pure scale-gauge quantity—rescaling *ρ*∞ does not change an exponential time constant—but it is model-conditional and practically sensitive, because it is inferred indirectly from the mutant overshoot together with unobserved initial amplitudes and target shifts. The gauge-invariant statements—a single nutrient-independent decay timescale, the nutrient-independent overshoot peak time, the monotonic rise of *ρ*∞ with nutrient, and the fourth-power dependence of the baseline-subtracted sensing signal on *ρ* —do not depend on this convention. Because the functionally important quantities coincide with the stiff, well-constrained directions, the memory lifetime and the nutrient-dependent acquisition rate we report are robust features of the fit.

### 2.6 Flagellar reprogramming runs near an information-geometric speed limit

The information-geometric speed limit bounds how fast the expectation of an observable can change along a differentiable path of probability distributions. For any observable *A*(*x*),

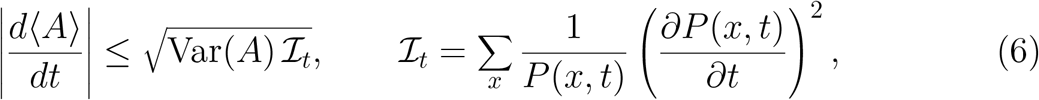

with the sum replaced by an integral for a continuous representation. Here *I*_*t*_ is the Fisher information of the distribution with respect to time, i.e. the squared statistical speed of the population through distribution space. Setting *A* = *x* flagella number, yields a model-free bound on the rate of change of the mean flagellar number *X*^*t*^ ≡ ⟨ *x* ⟩ . The quantities entering the bound—the variance and the time derivative of the histogram—are estimated directly from the time-resolved single-cell distributions, without using the fitted mean-field model.

Across the measured intervals, the observed rates | *dX*^*t*^*/dt* | lie below the Cramér– Rao bound and approach it during the periods of strongest reprogramming (Fig. 10). Because *∂*_*t*_*P* is estimated from a finite number of time points, the quantitative saturation ratio should be interpreted with the finite-difference and binning uncertainties reported in Methods; the robust conclusion is that the active phases operate close to, but not above, the model-free statistical-speed limit. In other words, during active remodeling *Salmonella* changes its mean flagellar number at close to the fastest rate compatible with its own single-cell variability and rate of distributional change. We stress the conditional content of this statement: the ceiling 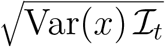 is set by the population’s own fluctuations rather than by an absolute biosynthetic maximum, and because the bound is kinematic, near-saturation reflects near-optimal use of the available statistical speed in the flagellar-number coordinate—not a thermodynamic or synthesis-rate maximum, and not evidence that fast remodeling is itself selected for. The information length, 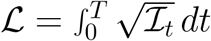, increases with nutrient in the wild type (ℒ = 1.03, 1.58, 2.15 at 0.2%, 0.5%, 1%), indicating greater distributional reprogramming in richer media. The mutant traverses comparable distances (1.42, 2.11) along a non-monotonic, out-and-back trajectory. This bound is kinematic: it follows from probability geometry and differentiability of *P* (*x, t*), not from thermodynamic dissipation. It is therefore complementary to the effective noise cost described above.

**Figure 10.**
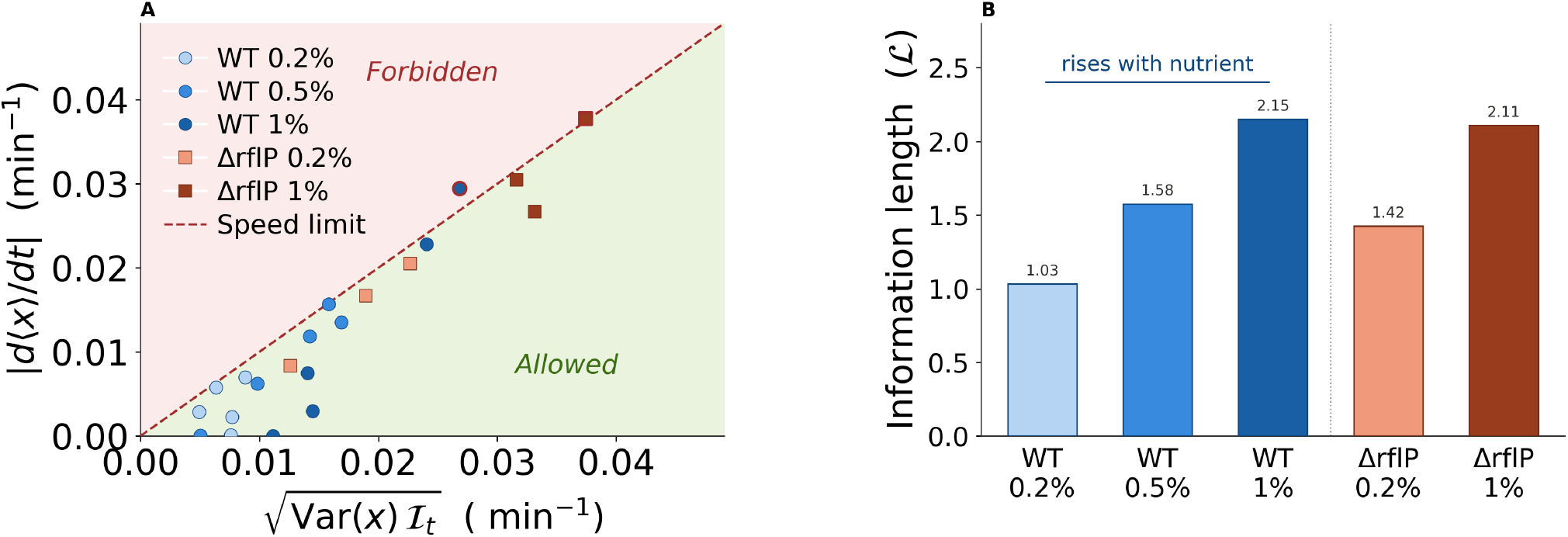
Flagellar reprogramming respects a model-free information-geometric speed limit. The measured rate | *d* ⟨ *x* ⟩ */dt* | versus the Cramér–Rao bound 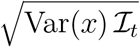 for the wild type and Δ*rflP* mutant; all observations lie on or below the bound (dashed diagonal), and active-phase points approach it. Inset/bar: the information length 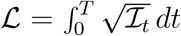 *dt* per condition, increasing with nutrient in the wild type.

## 3 Discussion

We developed a quantitative framework linking nutrient availability, flagellar production, and bacterial motility, and used it to dissect regulatory function kinetically. The model is built from two physically distinct layers on separated timescales: a fast RflP–FlhD_4_C_2_ regulatory layer that sets a correlation-dependent target *k*(*ρ*), and a slow physical layer in which synthesis, partition, and division track that target, so that flagellar number behaves as a leaky integrator of a latent sensing-memory variable. Using the Δ*rflP* mutant as a perturbation that removes the acquisition branch, we separated the nutrient-dependent acquisition of the regulatory correlation from its intrinsic decay, and showed that their ratio sets the steady-state fidelity, that precision is tuned through signal amplitude rather than integration time at a roughly twofold (∼ 1.7-fold) effective noise cost, and that the population reprograms flagellar number near a model-free information-geometric speed limit. A held-out nutrient condition confirms that the constrained quantities are predictive. The remainder of this section elaborates on what these findings mean and what bounds their interpretation.

The kinetic decomposition reframes how an RflP-type regulator should be understood. RflP is classically described as a repressor that, when active, lowers steady-state flagellar gene expression by targeting FlhD_4_C_2_ for degradation. Our analysis instead places it within a dynamical encoder: the functional quantity is not a static repression level but the *rate* at which the regulatory correlation is acquired, opposed by an intrinsic, RflP-independent decay revealed by the mutant overshoot. In this view, flagellar number is not a fixed, species-typical trait but a dynamically set readout shaped by recent nutrient history and by correlations inherited across division. The steep (fourth-power) dependence of the sensing signal on the steady-state correlation sharpens this point: because precision depends so strongly on the acquisition-to-decay balance, comparatively small kinetic changes—rather than large changes in mean expression—suffice to move the population between low-precision and high-precision regimes. This offers a concrete, measurable target for understanding how enteric bacteria graduate motility investment, and predicts that perturbations which change the acquisition rate (without altering decay) should reposition the population along the same fidelity axis.

The way precision is achieved departs from the classical picture of bacterial sensing precision. In the Berg–Purcell paradigm [25], a cell improves its estimate of an external concentration by integrating a noisy receptor signal over time, so that precision increases with the available averaging window—a bandwidth-limited strategy in which longer integration buys more independent samples. The encoding studied here does not operate this way: the relaxation time that would play the role of an integration window varies by less than a factor of two across nutrient conditions, while encoding precision varies by nearly two orders of magnitude. Mechanistically, the control variable is different—rather than averaging longer, the cell raises the steady-state regulatory correlation, and the steep dependence of the signal on that correlation converts modest kinetic changes into large precision changes. We stress that this is a contrast of strategy, not a violation of the Berg– Purcell bound, which concerns concentration estimation by receptor averaging and not the regulatory setting of flagellar number. The biological reading is that investing in signal amplitude is a route available specifically to an actively driven, far-from-equilibrium regulator: the same nonequilibrium driving that permits a strain-specific effective noise amplitude (the ∼ 1.7-fold cost) is what makes amplitude-tuning, rather than passive temporal averaging, the operative mechanism. The two physical limits we report are therefore complementary faces of one active process—a kinematic bound on the rate of distributional change and an effective cost of the precision it buys—and neither is a thermodynamic statement, since we measure no entropy production.

It is important to delineate what the data determine from what they do not. The analysis constrains the kinetic rates, the steady-state fidelity ratio, the effective precision cost, and the model-free speed bound; the parameter-identifiability analysis shows these to be the stiff, well-constrained combinations, and the held-out 0.4% condition confirms they are predictive. The data do *not* determine the absolute sensing correlation—only the gauge-invariant product *c* Γ(*ρ*) is identifiable, so the memory timescale carries a corresponding gauge dependence— nor the parametric form of the regulatory coupling, since over the fitted range the exact Γ(*ρ*) is close to a function quadratic at the origin, nor whether the cell literally performs Bayesian inference. We therefore treat the inference picture as the construction principle that fixes the form of the regulatory coupling, and confine the quantitative claims to the identifiable kinetic and information-theoretic quantities. This separation is a feature rather than a hedge: it isolates a small set of robust, falsifiable rate quantities from the modeling choices used to motivate them.

Some limitations need to be reported for the better understanding of the conclusions. They rest on a single experimental campaign with modest sampling: flagellar distributions were measured at a limited number of time points and nutrient concentrations, the wild type with three biological replicates and the Δ*rflP* mutant with two. The mutant overshoot reproduces across both replicates, but with only two replicates the replicate-based uncertainty remains coarse, and because the intrinsic decay timescale is extracted primarily from the mutant overshoot, its value should still be regarded as provisional; additional mutant replicates and finer sampling near the overshoot peak would tighten it substantially. The mutant growth rate was measured from OD_600_ and is comparable to the wild type at matched nutrient, which constrains the baseline division parameter; the residual uncertainty in the noise cost—which we accordingly report as a range rather than a point value—now stems mainly from the modest replicate number rather than from an unmeasured growth rate. The latent sensing correlation is never observed directly but inferred through the model, so the regulatory branch is tested only indirectly, through its kinetic and distributional consequences. The mean-field, linear-noise treatment is least reliable at low nutrient, where mean flagellar numbers are small and the discrete, non-Gaussian nature of the counts matters. Correspondingly, the information-geometric bound and the variance–relaxation relation are estimated from finite, binned histograms and carry sampling and discretization error, particularly in the sparsely populated tails. Finally, all measurements were obtained in well-mixed batch culture, so the spatial and temporal structure of natural nutrient environments is absent. These caveats constrain the quantitative claims but not the gauge-invariant qualitative conclusions: the acquisition–decay dichotomy between strains, a single intrinsic decay timescale, and a graded rather than switch-like nutrient response all survive them.

Several of these limitations point directly to next experiments. The most informative would make the latent variable observable: co-imaging a metabolic or regulatory reporter together with the flagellar reporter would convert the inferred sensing correlation into a measured quantity and provide a direct test of the two-layer construction. A second class of experiments would perturb the acquisition arm—through titratable *rflP* expression or the CsrA/ClpXP axis— which the model predicts should change the acquisition rate while leaving the intrinsic decay timescale unchanged, a sharp and falsifiable signature. Additional replicate-resolved mutant growth curves, together with denser sampling around the overshoot peak, would further tighten the estimates of *τ*_*ρ*_ and the effective noise-amplitude ratio. Beyond *Salmonella*, because nutrient fields are structured in space and time, extending the framework to spatial growth and to regulatory feedbacks beyond the single sensing channel would test how far the leaky-integrator picture generalizes; applying it to mixed communities, where interspecies signaling could shape motility investment, and to evolutionary dynamics, where the cost–precision trade-off could drive diversification, are natural longer-term directions.

In summary, by treating flagellar remodeling as a coupled regulatory and proliferative process, this study converts a static, species-typical description of flagellar number into a small set of measurable kinetic quantities—an acquisition rate, an intrinsic decay timescale, a fidelity ratio, an effective precision cost, and a model-free speed bound—and shows how their balance encodes nutrient information in bacterial flagellar heterogeneity.

## 4 Materials and methods

### 4.1 Bacterial strains and media

The *Salmonella enterica* subsp. *enterica* serovar Typhimurium strain LT2 used in this study was obtained from our lab collection. The strain EM3713 (*fliG*^22799^ mNeonGreen–FliG, N-terminal) harbored an mNeonGreen [10.1038/nmeth.2413] translational fusion on the N-terminus of the C-ring subunit FliG, which allowed direct visualization of the basal bodies, as they produced bright and consistent fluorescent foci that could be accurately visualized using epi-fluorescence microscopy and counted using automated image analysis. The bacteria were cultivated in M9 minimal medium supplemented with 0.2% glucose (w/v) and varying yeast extract concentrations (w/v: 0.2%, 0.4%, 0.5%, and 1%) aerobically at 37 °C with 180 rpm agitation. Overnight cultures were prepared by inoculating a single colony in M9 minimal medium supplemented with 0.2% glucose and 0.2% yeast extract. Day cultures were prepared by adjusting the OD_600_ of the overnight culture to 0.05 and grown in M9 minimal medium supplemented with 0.2% glucose and different yeast extract concentrations (0.2%, 0.4%, 0.5%, and 1%).

### 4.2 Growth rate measurements

The cultures were prepared as described above. Samples were taken at 0, 60, 90, 120, and 180 min after dilution of the overnight culture for OD_600_ measurements and subsequent microscopy imaging. An individual blank was used for each yeast extract concentration.

### 4.3 Fluorescence microscopy and image analysis

After measuring OD_600_, 2 *µ*L of the cells were spotted on a 1% agarose pad in PBS. The sample was air-dried for approximately 5 min (or until completely dry when needed), and a glass cover slide was carefully placed on top. Images were acquired using a Nikon Eclipse Ti2 inverted microscope, equipped with a CFI Plan Apochromat DM 60 × Lambda NA 1.40 Ph3 oil objective, an Orca Fusion BT camera (Hamamatsu) and a SPECTRA III LED light source (Lumencor). Images were captured with 500 ms exposure and 5% of the maximum illumination power. Z-stacks were taken in a 1 *µ*m range with three stacks of 0.5 *µ*m intervals. Images were analyzed using Fiji [26] with the MicrobeJ plugin [27].

### 4.4 Data analysis

For each cell, we counted the number of fluorescent mNeonGreen–FliG foci and used this count as the single-cell flagellar basal-body number, denoted *x*. Because cell-volume measurements were not used in the present analysis, *x* should be interpreted as a count per cell rather than as a true physical density. For the continuum model, the discrete count distribution was represented by a continuous approximation.

#### 4.4.1 Statistical tests

Goodness of fit was evaluated by comparing the model-predicted mean flagellar number with the experimental mean across time points. For each wild-type nutrient concentration and time point *t*, measurements were obtained from three independent biological replicates; let 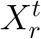 denote the empirical mean flagellar number in replicate *r*. The replicate-averaged mean, which estimates the population mean *X*^*t*^, is

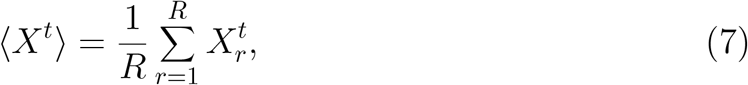

where *R* = 3, and the replicate standard deviation is

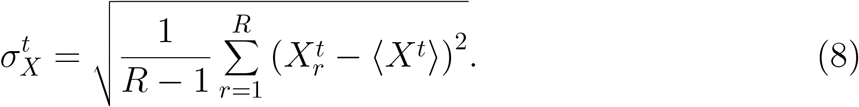

The model-predicted mean 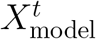 was compared with the replicate-averaged experimental mean ⟨*X*^*t*^⟩; agreement was summarized by

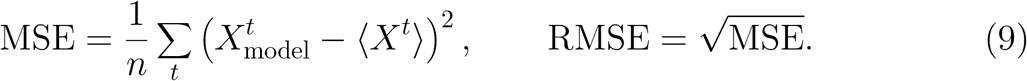

The held-out 0.4% wild-type condition is a prediction rather than a fitted condition, so its effective degrees of freedom should be interpreted separately from the fitted conditions. For the mutant conditions, two biological replicates are available; replicate-averaged means and replicate spreads are used, and because only two replicates contribute, MSE and RMSE are reported as the primary goodness-of-fit metrics rather than replicate-based *χ*^2^.

### 4.5 Regulatory inference and the two-layer construction

The model separates the cell into two layers operating on well-separated timescales. A fast **regulatory/biochemical layer** (the “software”) represents the RflP– FlhD_4_C_2_ signaling that computes how much flagellar synthesis the current environment warrants; biochemical signaling—reversible protein–protein binding and ClpXP-mediated proteolysis—updates this state continuously and rapidly. A slow **physical layer** (the “hardware”) is the number *X* of assembled flagella, which changes only through the discrete mechanics of synthesis and cell division. The flagellar count cannot change to satisfy a probability flux; it can only be synthesized and partitioned. We therefore apply the Bayesian sensing dynamics, adapted from our earlier work [19], strictly to the regulatory layer, whose output sets the synthesis target that the physical layer then tracks. The Bayesian channel enters as a *premise* that supplies the functional form of the regulatory coupling; it is not a claim that these data prove explicit Bayesian inference by the cell.

#### The sensing chain

We model the transduction of the nutrient signal as the linear chain (as shown in Fig. 1)

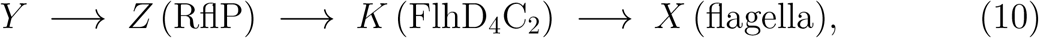

where the nutrient condition *Y* is transduced into an RflP-dependent regulatory state *Z*, which sets the activity *K* = FlhD_4_C_2_ of the master regulator, which in turn drives flagellar synthesis *X*. We stress that *Z* is an effective regulatory coordinate representing the nutrient-dependent state of the RflP–FlhD_4_C_2_ branch, *not* RflP abundance or activity itself: because RflP represses FlhD_4_C_2_, increasing nutrient *relieves* this repression, and we orient *Z* so that larger *Z* corresponds to reduced repression and increased flagellar regulatory activity. Each stage is modeled as linear–Gaussian. The Bayesian update operates on the *Z* → *K* step (the regulatory state informing the master regulator), and *Z* is the sufficient statistic linking *K* to *Y* : conditioned on *Z*, the regulator *K* is independent of the input *Y* (the Markov property of the chain). Under the linear–Gaussian and mean-state closure used below, the regulatory drive is parameterized by the internal sensing–regulation correlation

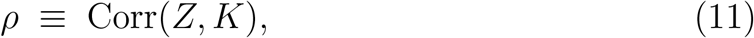

the correlation between the RflP-dependent regulatory state and the master-regulator activity, which we take as the dynamical sensing variable throughout. We emphasize that *ρ* is a *latent* quantity: it is an internal regulatory correlation that we do not measure directly, distinct from the experimentally accessible nutrient–flagella correlation Corr(*X, Y* ). As shown below, *ρ* enters the dynamics only through the product *c* Γ(*ρ*), so its absolute value is not identifiable from the data (Section 4.10). In the Δ*rflP* mutant, where RflP is deleted, *ρ*(*t*) should be read as an effective inherited regulatory-memory coordinate projected onto the same model variable, rather than as a literal correlation with an RflP readout; the mutant therefore isolates the *decay* of this inherited effective state rather than a measurable RflP-state correlation.

#### The regulatory layer (Bayesian update of *K*)

The cell maintains an internal distribution *P*_reg_(*K, t*) over the regulatory activity and updates it toward the Bayesian posterior given the sensor readout *Z*, over a fast biochemical relaxation time *τ*_*B*_,

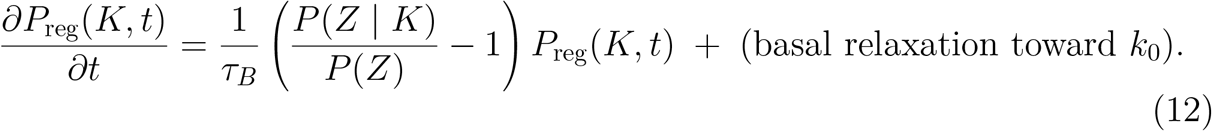

Because *Z* is the sufficient statistic for *K* along the chain (10) (conditioned on *Z*, the regulator *K* is independent of the nutrient input *Y* ), the update can be written directly in terms of the sensor *Z* rather than the raw nutrient *Y* . The derivation of Eq. (12) from the one-step Bayesian update is given in [19] and is not reproduced here. The relaxation time *τ*_*B*_ is physically distinct from the correlation-decay time *τ*_*ρ*_ introduced in Section 2.3: the former sets the rate of the inference update, the latter the rate at which the internal correlation *ρ* is lost in the absence of acquisition.

#### Collapse to the coupling Γ(*ρ*)

Because the chain (10) is linear–Gaussian and *Z* is the sufficient statistic, the likelihood ratio depends on the regulatory state only through *ρ* = Corr(*Z, K*), and can be written in terms of the standardized sensor variable and the correlation parameter,

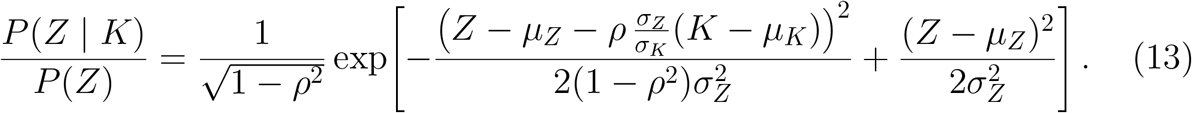

To close the dynamics at the population level we adopt a mean-field closure: the regulatory drive of the population is evaluated at its mean state, i.e. at *Z* = *µ*_*Z*_ and *K* = *µ*_*K*_, for which the exponent in Eq. (13) vanishes identically. The bracketed factor in Eq. (12) then reduces, at the mean state, to

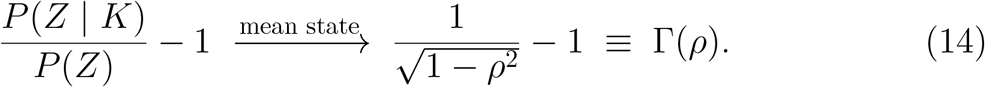

This mean-state evaluation, rather than a full population average, is the operative closure: averaging the bare likelihood ratio over the internal distribution returns unity by normalization (E_*P*(*K*)_[*P* (*Z K*)*/P* (*Z*)] = 1), so the regulatory drive Γ(*ρ*) is specifically the drive of the mean (typical) cell. The exact regulatory coupling is 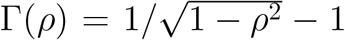. For analytic tractability, the closed-form mean-field trajectories used in all reported fits employ its small-correlation expansion,

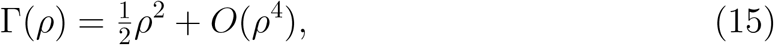

which also underlies the fourth-power signal scaling below; numerical integration of the mean-field equations is overlaid in Fig. 5 as a consistency check on these closed forms. We note that the fitted steady-state correlations are not uniformly small (*ρ*_∞_ ranges up to ≈ 0.60 at high nutrient; Section 2.3.2, Table 2), so at the upper end Γ(*ρ*) departs from *ρ*^2^*/*2 by up to ∼ 25%. However the fitting metrics show a good agreement with experimental data (Table 3). This might happen because only the gauge-invariant product *c* Γ(*ρ*)—equivalently the sensing-dependent target shift *k*(*ρ*) − *k*_0_—is matched to the data (Section 4.10), this expansion enters as part of the latent-correlation parameterization rather. The numerical correlation values at high nutrient should be read accordingly.

#### Quasi-steady regulatory target

Taking the mean of Eq. (12) under the mean-state closure, the mean regulatory activity relaxes toward a baseline *k*_0_ shifted upward by the regulatory drive *c* Γ(*ρ*), where *c* is the coupling of the sensing drive to the promoter,

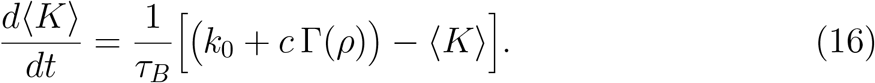

Here *k*_0_ is the blind, nutrient-free baseline toward which *K* relaxes in the absence of an informative signal, and *c* Γ(*ρ*) is the regulatory drive that raises the target when the internal readout is correlated with the environment. The regulatory relaxation time *τ*_*B*_ is set by the turnover of the FlhD_4_C_2_ complex through ClpXP-mediated proteolysis, the mechanism that exists precisely to enable rapid flagellar responses to environmental change [14]. The measured half-life of the FlhD_4_C_2_ complex is approximately 2–3 min [28]; while *τ*_*B*_ is not identical to this half-life—it also includes the upstream RflP binding step—it is of the same order, *τ*_*B*_ ∼ *O* (a few minutes). This is an order of magnitude shorter than the timescales governing the physical layer: the flagellar relaxation time *τ*_*X*_ = 1*/*(*µ*_0_ − *ak*) ≈ 43–60 min (Section 4.10) and the sensing-memory timescale *τ*_*ρ*_ ≈ 400 min (Section 2.3.1), giving the ordering

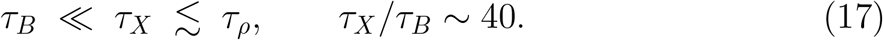

On the timescale of flagellar assembly the regulatory network is therefore always at quasi-steady state. Setting *d* ⟨ *K* ⟩ */dt* = 0 in Eq. (16), the fast regulatory layer is adiabatically slaved to its target,

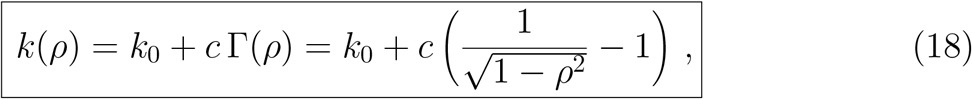

evaluated at the current, slowly-varying correlation *ρ*(*t*). The correlation *ρ*(*t*) is *not* eliminated: it is the slow sensing-memory variable, with its own leaky-integrator dynamics (Section 2.3); only the fast biochemical relaxation of *K* toward *k*(*ρ*) is adiabatically removed. Because *ρ* enters only through *c* Γ(*ρ*), the pair (*c, ρ*) is fixed by the data only up to a gauge transformation; we fix *c* and report gauge-invariant quantities (Section 4.10). We do not claim *k*(*ρ*) is an optimal target, only that it is the Bayesian-motivated regulatory drive evaluated at the mean state.

#### Two routes from nutrient to flagella

Nutrient influences flagellar number through two routes: a regulatory route (*Y* → *Z → K* → *X*), which enters through Γ(*ρ*) and requires a functional RflP–FlhD_4_C_2_ branch, and a proliferation route (*Y* → *µ*_0_ → *X*), through the nutrient-dependent division/growth parameter *µ*_0_(*Y* ) (Section 4.6). In the wild type the regulatory route drives the acquisition of the internal correlation *ρ*. In the Δ*rflP* mutant the RflP-dependent acquisition term is severed (no acquisition source, *r*_WT_ = 0), so *ρ* represents only an inherited correlation that decays with no replenishment. We caution, however, that the mutant is not a pure physical-channel-only system: the fitted baseline target *k*_0_ retains a substantial nutrient dependence (Table 3, *k*_0_ rising from ≈ 0.2054 at 0.2% to ≈ 2.0172 at 1%), indicating that nutrient dependence still enters through RflP-independent effects on basal synthesis and proliferation. We therefore interpret the Δ*rflP* mutant primarily as isolating the *decay* of the inherited effective regulatory state, rather than as a system in which all nutrient response has been removed.

### 4.6 The physical layer: proliferation dynamics of flagellar number

The regulatory target *k*(*ρ*) enters the physical dynamics as the synthesis set-point that the flagellar count tracks through growth and division. We derive the evolution of the flagellar distribution from the underlying birth–partition process.

#### Synthesis and partition

Over a cell cycle a mother cell accumulates new flagella through the FlhD_4_C_2_–FliA hierarchy at a rate set by the regulatory target *k*(*ρ*). Modeling synthesis as a Poisson process with mean *k*(*ρ*),

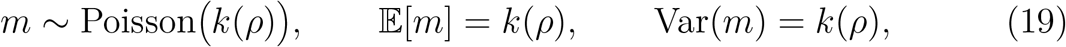

so that the pre-division count is *z* = *x*^′^ + *m*. At division each flagellum is inherited by a given daughter independently with probability 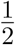,

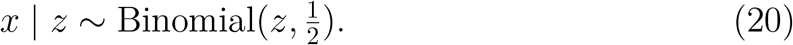

Combining Poisson synthesis with binomial partition, the first two jump moments of the transition from a mother with *x*^′^ flagella are

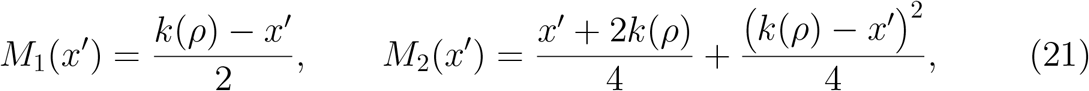

the drift *M*_1_ pulling the count halfway toward the target at each division and *M*_2_ collecting the Poisson and binomial variances.

#### Load-dependent division

Division is slowed by flagellar load, reflecting competition of flagellar biosynthesis with the growth machinery for ribosomes and energy,

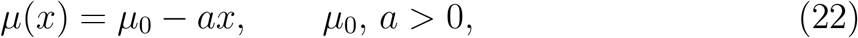

where *µ*_0_ is the baseline proliferation rate (measured from OD_600_ growth curves at each nutrient concentration) and *a* the per-flagellum biosynthetic cost (fixed at *a* = 2.65 × 10^−4^); the regime *aX*^*t*^ ≪ *µ*_0_ holds *a posteriori* in all conditions.

#### Master equation and Kramers–Moyal expansion

Let *n*(*x, t*) be the number of cells with *x* flagella. Cells leave state *x* by dividing at rate *µ*(*x*) and enter as daughters of cells in other states, with both daughters distributed by the inheritance kernel *W* (· | *x*^′^); the factor of two counts the two daughters,

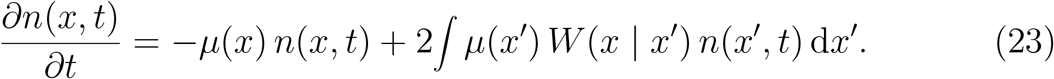

Expanding the inheritance integral to second order in the jump size (Kramers– Moyal truncation, justified when typical jumps are small relative to typical *x*) gives

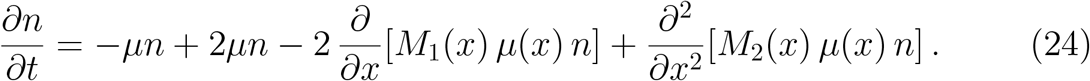

The first two terms combine to a net birth +*µ*(*x*) *n*, and the factor of two from daughter doubling exactly cancels the factor of 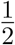 in *M*_1_ = (*k*(*ρ*) − *x*)*/*2, leaving a clean drift toward the target. Defining the effective diffusion coefficient consistently with the second-order term of Eq. (24),

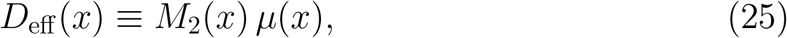

and normalizing to the probability *P* (*x, t*) = *n*(*x, t*)*/N* (*t*) with *N* (*t*) = ∫ *n dx* total population grows at the mean division rate,

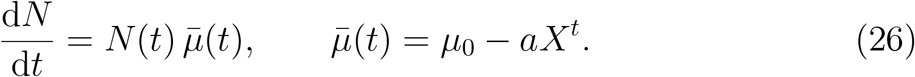

Applying the quotient rule to *P* = *n/N* yields the evolution equation for the flagellar distribution,

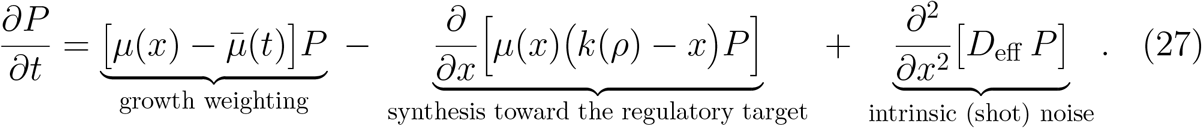

The growth-weighting term reweights lineages by their relative division rate and conserves total probability after normalization; the drift relaxes the distribution toward the regulatory target *k*(*ρ*); and *D*_eff_ collects the Poisson synthesis and binomial partition noise. Crucially, the Bayesian sensing dynamics enter Eq. (27) *only* through the target *k*(*ρ*) in the drift: the inference sets the set-point, and the physical layer tracks it. No likelihood-ratio term acts on *P* (*x, t*) directly, so flagellar number changes only through synthesis and division, as it must.

##### Mean-field reduction

Taking the first moment of Eq. (27) and closing at the mean (Section 4.5) gives the mean-field equation for the population mean flagellar number *X*^*t*^ ≡ ⟨*x*⟩,

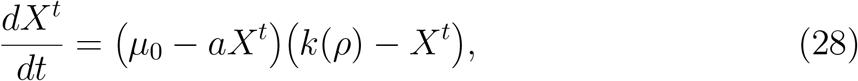

which relaxes toward the regulatory target *k*(*ρ*) on the timescale *τ*_*X*_ = 1*/*(*µ*_0_ − *ak*). The target itself moves on the slow sensing-memory timescale through *ρ*(*t*), so the observed dynamics are the physical layer chasing a slowly drifting set-point: building when *ρ* rises in the wild type, and relaxing back toward *k*_0_ when *ρ* decays in the Δ*rflP* mutant (Section 2.3).

##### Flagellar number and effective density

Experimentally, the measured single-cell observable is the number of foci in each bacterium, which reports the number of flagellar basal bodies per cell. We denote this discrete count by *n*. In the stochastic and continuum descriptions below, we use the variable 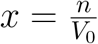, where (*V*_0_) is the bacterial cell volume. Because individual cell volumes were not measured in the present analysis, and because our model tracks flagellar-state variation per cell rather than physical concentration in (*µ*m^−3^), we set (*V*_0_ = 1) for simplicity. Thus *x* is numerically identical to the basal-body count (*n*), but is written as an effective flagellar density in the continuum master-equation, Fokker–Planck, and mean-field formulations.

Accordingly, throughout the mathematical model, flagellar density refers to a normalized per-cell state variable, or flagellar number per unit reference cell volume, not to an independently measured volumetric density. All model quantities such as (*P* (*x, t*)), (*X*_*t*_ = ⟨*x*⟩), and the regulatory target (*k*(*ρ*)) can therefore be read equivalently as distributions, means, and targets of basal-body number per cell under the unit-volume normalization.

### 4.7 Analytical approximation for the mean-field trajectories

The full mean-field equation,

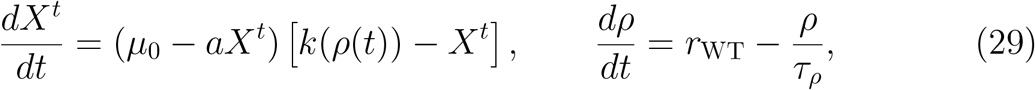

is nonlinear because the relaxation rate depends on *X*^*t*^. The closed-form expressions used for intuition and fitting are therefore best described as a linearized analytical approximation, not as the exact solution of the full nonlinear ODE. The correlation dynamics are exactly

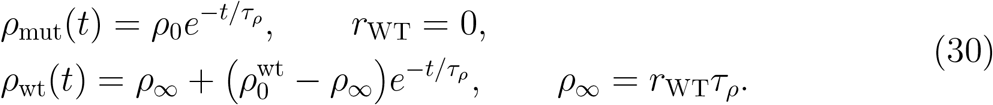

Throughout the closed-form trajectories we use the small-correlation expansion

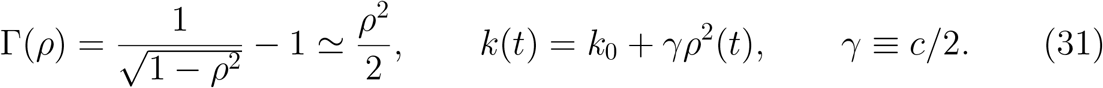

As noted in Section 4.5, the fitted *ρ*∞ reach ≈ 0.60 at high nutrient, beyond the nominal small-*ρ* regime, so at the upper end this expansion carries an ∼ 25% departure in Γ; the reported high-nutrient *ρ*∞ are therefore fixed-*c* gauge-convention estimates rather than calibrated correlations, while the gauge-invariant kinetic conclusions are unaffected. The remaining approximation is to linearize the flagellar dynamics around the relevant operating point, replacing the state-dependent relaxation rate by a constant rate. This yields sums of exponentials that expose the time scales of the response.

#### Mutant Δ*rflP* trajectory

For the mutant, 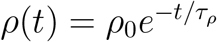, so the target inherits twice this decay rate through 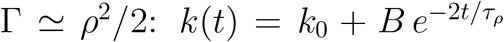 with 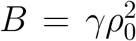.

Linearizing around the baseline *k*_0_ gives the relaxation rate

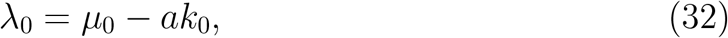

and the trajectory

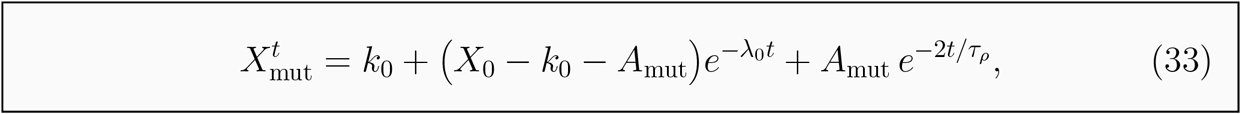

where

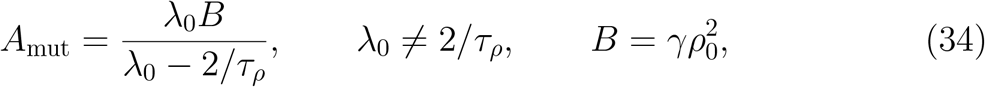

and *γ* = *c/*2 is the small-correlation coefficient of the target expansion (see below). The two exponentials respectively describe relaxation of the physical flagellar count and decay of the inherited sensing-dependent boost.

#### Peak time for Δ*rflP*

Setting *dX*^*t*^*/dt* = 0 in Eq. (33) gives

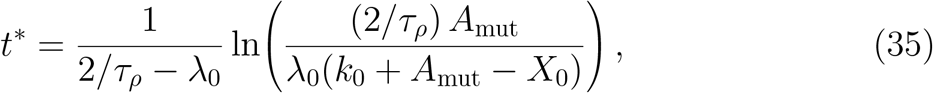

when the argument of the logarithm is positive and greater than one. Thus a finite overshoot requires the initial sensing-dependent boost to be large enough to drive *X*^*t*^ above its long-time baseline.

#### Wild-type trajectory

For the wild type, 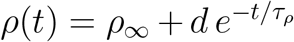 with 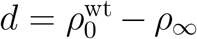. The target can be written

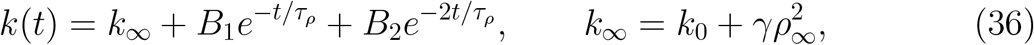

where *B*_1_ = 2*γρ*_∞_ *d* and *B*_2_ = *γ d*^2^. Linearizing around *k*_∞_ gives *λ*_∞_ = *µ*_0_ − *ak*_∞_ and

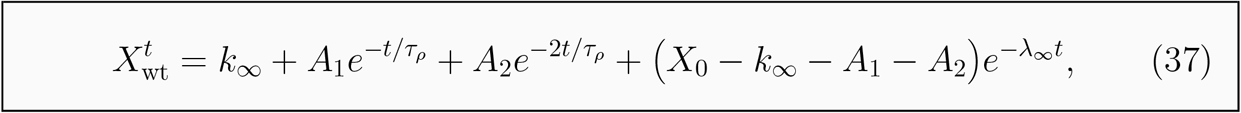

with

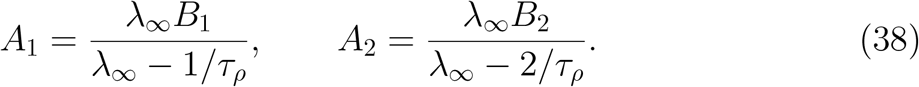

The long-time sensing-dependent increase in target is 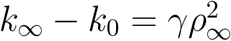; the fourth-power precision scaling discussed in the Results follows only after converting this excess target into signal power.

### 4.8 SNR calculation

We define the signal-to-noise ratio in the transformed variable as

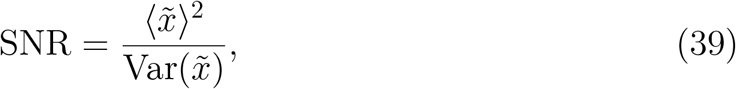

where 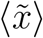 and 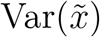 are the mean and variance of the log-transformed count 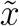 for a given condition and time point. Writing *δX* = *X*^*t*^ − *k*(*ρ*) for the fluctuation about the operating point and linearizing Eq. (29) gives

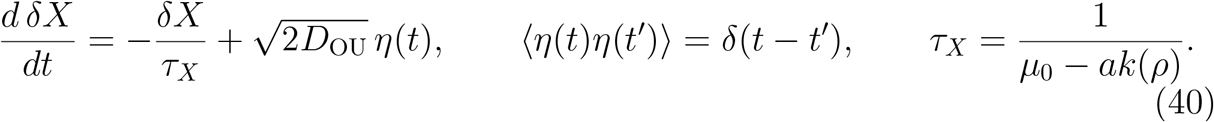

The stationary variance of this Ornstein–Uhlenbeck approximation is 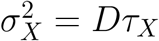 In the log-transformed analysis, the same relation is used with the empirical variance of 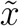 to estimate an effective noise amplitude 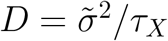. Because the system is growing and actively regulated, *D* is an effective nonequilibrium noise amplitude, not a thermodynamic temperature.

### 4.9 Information-geometric speed limit

For each condition, the empirical distributions *P* (*x, t*) were estimated using a fixed histogram binning across time points. The time derivative *∂*_*t*_*P* (*x, t*) was estimated by finite differences between adjacent time points. The Fisher information with respect to time was computed as

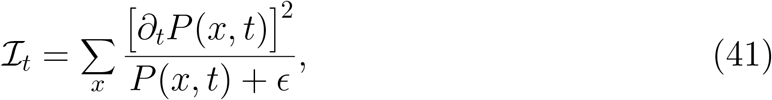

where *ϵ* is a small pseudocount used only to avoid division by empty bins. The information length was approximated by trapezoidal integration of 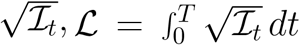. We summarize how closely the bound is approached by the saturation ratio

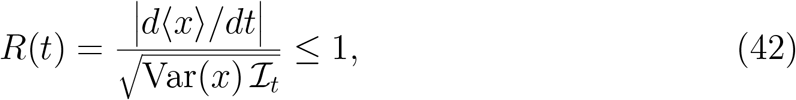

which is unity at the Cramér–Rao bound. All quantities entering Eqs. (41)–(42) are computed directly from the time-resolved single-cell histograms, without the fitted mean-field model. Uncertainty was estimated by bootstrap resampling of cells within replicates and repeating the full histogram and finite-difference procedure; the finite number of time points and the binning of sparsely populated tails are the dominant sources of error.

### 4.10 Parameter estimation and fitting strategy

#### Parameter inventory

The mean-field model (29) together with the correlation dynamics (3) contains parameters of three kinds. *Fixed* constants: the biosynthetic cost *a* = 2.65 × 10^−4^, taken from the literature on flagellar biosynthetic burden in *Salmonella*, and the sensing coupling *c*, which is fixed to a reference value (*c* = 100) because only the gauge-invariant product *c* Γ(*ρ*) is identifiable (Section 4.10). *Shared* parameters, common to all conditions of a strain: the intrinsic correlation-decay time *τ*_*ρ*_ (shared across both strains, as it represents a nutrient-independent forgetting process), and the initial conditions 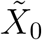 and *ρ*_0_. *Per-condition* parameters: the baseline division/growth parameter *µ*_0_(*Y* ) (measured independently from the OD_600_ growth curves at each nutrient concentration, for both the wild type and the Δ*rflP* mutant; the two strains grow at comparable rates at matched nutrient, with *µ*_0_ differing by less than ∼ 3%), the baseline target *k*_0_(*Y* ), and—for the wild type only—the acquisition rate *r*_WT_(*Y* ), with *r*_WT_ = 0 enforced for the mutant.

#### Two-stage estimation

Because the mutant has no acquisition source (*r*_WT_ = 0), its dynamics depend on *τ*_*ρ*_ but not on any wild-type-specific rate; the mutant overshoot therefore isolates the shared decay time. We exploit this with a two-stage procedure. *Stage 1:* the two mutant nutrient conditions are fit jointly for the shared *τ*_*ρ*_, the shared mutant initial conditions 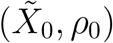, and the per-condition baseline target *k*_0_(*Y* ), with the division/growth parameter *µ*_0_(*Y* ) fixed at its OD_600_-measured value (above). The fit minimizes the summed squared deviation between the analytical mean-field trajectory and the replicate-averaged measured mean flagellar number. Replicate spreads are used for plotting uncertainty and goodness-of-fit diagnostics. relaxation of the overshoot determine *τ*_*ρ*_ (Section 2.3.1). *Stage 2:* with *τ*_*ρ*_ fixed at its Stage-1 value, each wild-type condition is fit for its (*k*_0_, *r*_WT_), holding the wild-type initial conditions at their shared values. This ordering reflects the causal structure of the model: the knockout fixes the intrinsic timescale, and the wild-type response then fixes the nutrient-dependent acquisition.

#### Optimization

The reported parameter fits minimize the unweighted mean-squared deviation between the analytical mean-field trajectory (*X*_model_(*t*)) and the replicate-averaged experimental mean ⟨*X*_*t*_⟩,

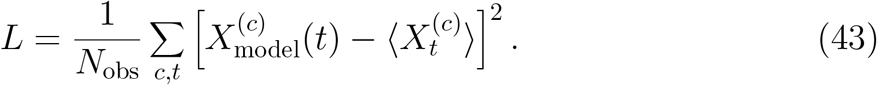

Replicate standard deviations are used for plotting uncertainty and goodness-of-fit diagnostics.

#### Confidence intervals

Uncertainties on the kinetic parameters are obtained by profile likelihood. For each parameter of interest the parameter is fixed on a grid while all remaining parameters are re-optimized, tracing the profile *Dχ*^2^; the 68% and 95% confidence intervals correspond to *Dχ*^2^ ≤ 1 and ≤ 3.84, respectively (one degree of freedom). The resulting interval on the acquisition rate *r*_WT_(*Y* ) is propagated, together with the interval on *τ*_*ρ*_, to the steady-state encoding precision by Monte Carlo sampling, yielding the uncertainty band on SNR∞ . We note that the profile interval on *τ*_*ρ*_ is a within-convention statistical interval; the dominant uncertainty on the memory lifetime *τ*_*ρ*_ is its model-conditional, practical-identifiability sensitivity (Section 4.10), so we report *τ*_*ρ*_ ≈ 400 min under the reference parameterization rather than the narrow profile interval alone.

#### Out-of-sample validation

The 0.4% wild-type condition is withheld from the primary predictive fit. Its parameters (*µ*_0_, *k*_0_, *r*_WT_) are predicted by interpolating the fitted nutrient dependences from the remaining conditions, and the predicted mean trajectory and steady-state precision are compared to the held-out measurements (Section 2.3.2). Please note that, the 0.4% WT condition is withheld from the primary model fit and is used as an interpolation test of the nutrient-dependent trends. In a separate post-hoc diagnostic, we also fitted the 0.4% condition to examine how it would enter the Fisher-information structure. This diagnostic fit is not used to claim out-of-sample prediction.

### Identifiability and sloppiness analysis

To establish that the biological conclusions rest on well-determined quantities, and to characterize which parameter combinations the data constrain, we analyzed the curvature of the fit objective at the best-fit point. For this diagnostic we use the curvature of the fitted objective, rather than bootstrap resampling, because the FIM is designed to reveal local parameter degeneracies.

#### Cost landscape

We first map the fit objective (43) over pairs of parameters about the best fit (Fig. 11). Elongated, curved valleys in these landscapes indicate parameter combinations that the data constrain only weakly, while sharply curved directions indicate well-determined combinations.

**Figure 11.**
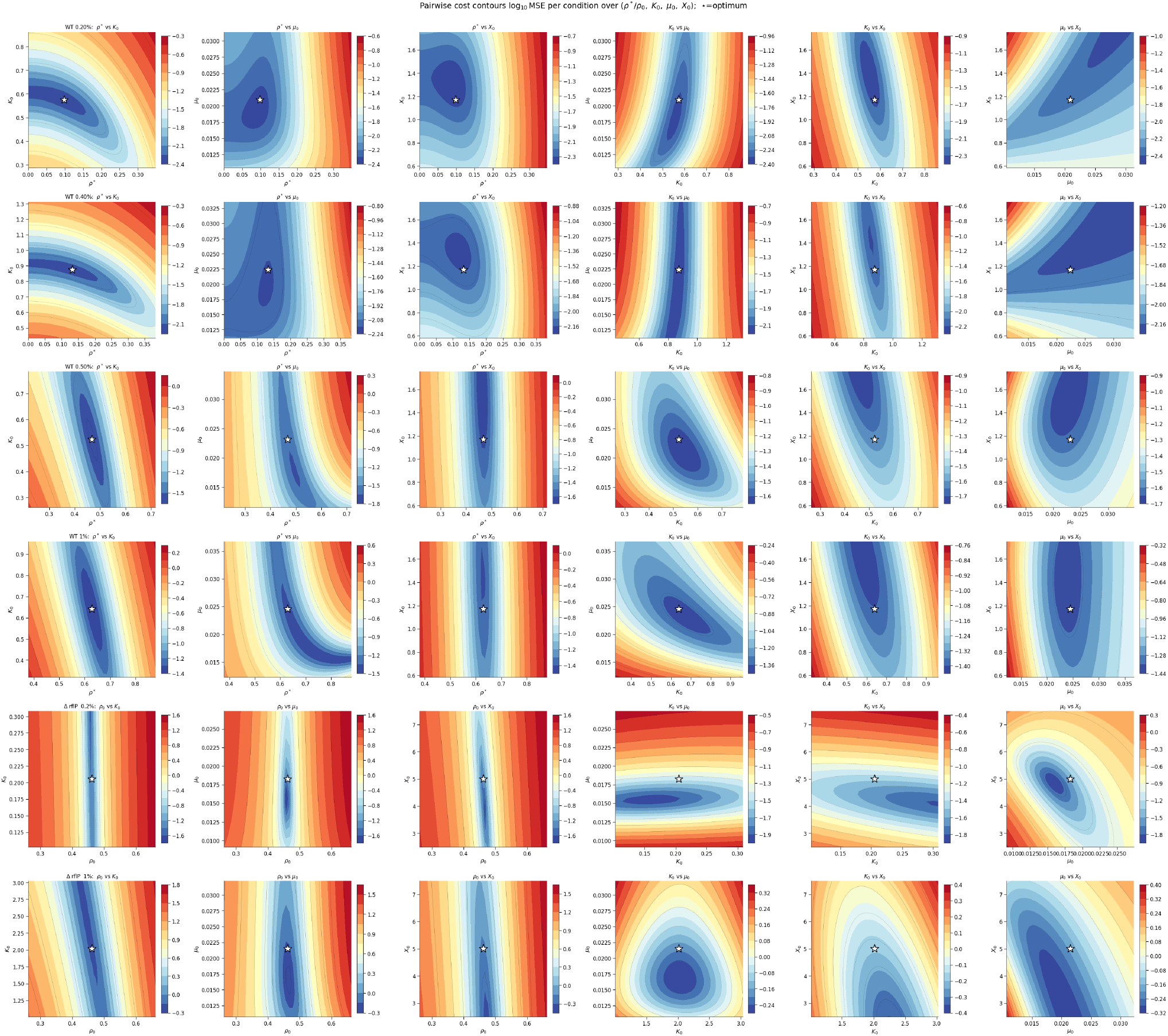
Identifiability analysis for 3 parameters for each 3 concentrations of WT strain and Δ*rflP* mutant strain at 2 concentrations.

#### Fisher-information eigenspectrum

We quantify this with the Fisher information matrix (FIM), evaluated at the best fit in log-parameter space,

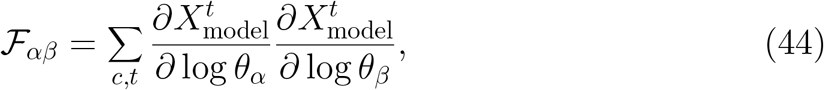

whose eigenvalues measure how stiffly the objective rises along the corresponding eigenvector directions. The sum runs over all nutrient conditions *c* and time points *t* for both strains, so *F* is a single global matrix over the full set of 16 model parameters (the shared decay time and initial conditions together with the per-condition baseline targets and acquisition rates), and the eigenspectrum reported in Fig. 9 is that of this global matrix rather than a collection of per-condition spectra. The spectrum spans several orders of magnitude—the hallmark of a “sloppy” model, in which a few stiff combinations dominate the fit while the remainder are weakly constrained.

#### Stiff directions are functional; sloppy directions are gauge

The structure of the eigenvectors is informative rather than arbitrary. The stiffest directions load on the intrinsic decay rate 1*/τ*_*ρ*_ (together with the mutant overshoot amplitude) and on the wild-type acquisition rates *r*_WT_(*Y* )—equivalently, on the steady-state correlation *ρ*_∞_ = *r*_WT_*τ*_*ρ*_, which sets the encoding fidelity. The sensing coupling *c* is fixed at the reference value *c* = 100 and is therefore not a parameter of this FIM; the structural *c*–*ρ* gauge—under which 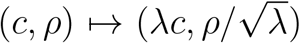 leaves the product *c* Γ(*ρ*) invariant to leading order—is exposed analytically and is what motivates fixing *c*. Within the reference gauge, the sloppiest FIM directions load on the latent-correlation amplitudes (the initial correlation *ρ*_0_ and the per-condition steady-state correlations) and on the unobserved initial condition 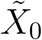. Thus the data determine precisely the combinations the theory identifies as functional (the kinetic rates and the fidelity ratio) and leave unconstrained precisely the latent-amplitude and initial-state directions.

#### Identifiability of the memory lifetime

The absolute scale of the latent correlation *ρ* and the acquisition rates *r*_WT_ are gauge-dependent, since only the product *c* Γ(*ρ*) enters the dynamics; quantities sensitive to this scale should be reported cautiously. The memory lifetime *τ*_*ρ*_ is not a pure scale-gauge quantity—rescaling *ρ* does not by itself change an exponential time constant—but its numerical estimate is model-conditional and practically sensitive, because it is inferred indirectly from the mutant overshoot together with the unobserved initial amplitudes and target shifts, and different parameterizations that fit the data comparably yield somewhat different values. We therefore report *τ*_*ρ*_ ≈ 400 min under the reference parameterization and emphasize that the gauge-invariant statements—the existence of a single nutrient-independent decay timescale, the nutrient-independence of the overshoot peak time, the monotonic rise of *ρ*∞ with nutrient, and the fourth-power dependence of the baseline-subtracted sensing signal on *ρ*∞ —do not depend on this convention. Co-imaging of the RflP readout with flagellar number would make the latent correlation observable, fix its absolute scale, and remove the residual parameterization sensitivity in *τ*_*ρ*_.

#### Functional sensitivity

Finally, we computed the sensitivity of the steady-state encoding precision SNR_∞_ ∝ Γ(*ρ*∞ )^2^ to each parameter, |*∂* log SNR_∞_ */∂* log *θ*_*α*_ |. The sensitivity is concentrated on (*r*_WT_) and (*τ*_*ρ*_) because (*ρ*∞ = *r*_WT_*τ*_*ρ*_), their log-sensitivities have the same sign. Equivalently, the sensitivities to (*r*_WT_) and to the decay rate (1*/τ*_*ρ*_) have opposite signs and is negligible along the gauge and initial-condition directions, confirming that the functionally relevant output is controlled by the same stiff combinations that the data constrain.

## Acknowledgments and funding

AB thanks Khalifa University for the support. HH, ME and MG would like to thank Volkswagenstiftung for its support of the “Life?” program (96732). Finally, HH acknowledges the support of the RIG-2025-001 grant from Khalifa University and the UAE-NIH Collaborative Research grant AJF-NIH-25-KU.

## Conflicts of interest

The authors have no conflicts of interest.

## S.I. Supplementary Information

## Notes

### Competing Interest Statement

The authors have declared no competing interest.

